# MEArec: a fast and customizable testbench simulator for ground-truth extracellular spiking activity

**DOI:** 10.1101/691642

**Authors:** Alessio P. Buccino, Gaute T. Einevoll

## Abstract

When recording neural activity from extracellular electrodes, both *in vivo* and *in vitro*, spike sorting is a required and very important processing step that allows for identification of single neurons’ activity. Spike sorting is a complex algorithmic procedure, and in recent years many groups have attempted to tackle this problem, resulting in numerous methods and software packages. However, validation of spike sorting techniques is complicated. It is an inherently unsupervised problem and it is hard to find universal metrics to evaluate performance. Simultaneous recordings that combine extracellular and patch-clamp or juxtacellular techniques can provide ground-truth data to evaluate spike sorting methods. However, their utility is limited by the fact that only a few cells can be measured at the same time. Simulated ground-truth recordings can provide a powerful alternative mean to rank the performance of spike sorters. We present here `MEArec`, a Python-based software which permits flexible and fast simulation of extracellular recordings. `MEArec` allows users to generate extracellular signals on various customizable electrode designs and can replicate various problematic aspects for spike sorting, such as bursting, spatio-temporal overlapping events, and drifts. We expect `MEArec` will provide a common testbench for spike sorting development and evaluation, in which spike sorting developers can rapidly generate and evaluate the performance of their algorithms.

## 1 Introduction

Extracellular neural electrophysiology is one of the most used and important techniques to study brain function. It consists of measuring the electrical activity of neurons from electrodes in the extracellular space, that pick up the electrical activity of surrounding neurons. To communicate with each other, neurons generate action potentials, which can be identified in the recorded signals as fast potential transients called *spikes*.

Since electrodes can record the extracellular activity of several surrounding neurons, a processing step called spike sorting is needed. Historically this has required manual curation of the data, which in addition to being time consuming also introduces human bias to data interpretations. In recent years, several automated spike sorters have been developed to alleviate this problems. Spike sorting algorithms [43, 29] attempt to separate spike trains of different neurons (units) from the extracellular mixture of signals using a variety of different approaches. After a pre-processing step that usually involves high-pass filtering and re-referencing of the signals to reduce noise, some algorithms first detect putative spikes above a detection threshold and then cluster the extracted and aligned waveforms in a lower-dimensional space [41, 44, 9, 23, 26]. Another approach consists of finding spike templates, using clustering methods, and then matching the templates recursively to the recordings to find when a certain spike has occurred. The general term for these approaches is template-matching [40, 47, 11]. Other approaches have been explored, including the use of independent component analysis [25, 3] and semi-supervised approaches [28].

The recent development of high-density silicon probes both for *in vitro* [2, 14] and *in vivo* applications [37, 27] poses new challenges for spike sorting [45]. The high electrode count calls for fully automatic spike sorting algorithms, as the process of manually curating hundreds or thousands of channels becomes more time consuming and less manageable. Therefore, spike sorting algorithms need to be be capable of dealing with a large number of units and dense probes. To address these requirements, the latest developments in spike sorting software have attempted to make algorithms scalable and hardware-accelerated [40, 26, 47, 39].

The evaluation of spike sorting performance is also not trivial. Spike sorting is unsupervised by definition, as the recorded signals are only measured extracellularly with no knowledge of the underlying spiking activity. A few attempts to provide ground-truth datasets, for example by combining extracellular and patch-clamp or juxtacellular recordings [22, 20, 37, 47, 33, 1] exist, but the main limitation of this approach is that only one or a few cells can be patched at the same time, providing very limited ground-truth information with respect to the number of neurons that can be recorded simultaneously from extracellular probes. An alternative method consists of adding artificial or previously-sorted and well-isolated spikes in the recordings (hybrid method) [44, 46]. The hybrid approach is convenient as all the characteristics of the underlying recording are kept. However, only a few hybrid units can be added at a time, and this limits the validation capability of this method.

Biophysically detailed simulated data provide a powerful alternative and complementary approach to spike sorting validation [12]. In simulations, recordings can be built from known ground-truth data for all neurons, which allows one to precisely evaluate the performance of spike sorters. Simulators of extracellular activity should be able to replicate important aspects of spiking activity that can be challenging for spike sorting algorithms, including bursting modulation, spatio-temporal overlap of spikes, unit drifts over time, as well as realistic noise models. Moreover, they should allow users to have full control over these features and they should be efficient and fast. While simulated recordings provide ground-truth information of many units at once, it is an open question how realistically they can reproduce real recordings.

In the last years, there have been a few projects aim to develop neural simulators for benchmarking spike sorting methods [7, 19, 35]: Camunas et al. developed `NeuroCube` [7], a MATLAB-based simulator which combines biophysically detailed cell models and synthetic spike trains to simulate the activity of neurons close to a recording probe, while noise is simulated by the activity of distant neurons. `NeuroCube` is very easy to use with a simple and intuitive graphical user interface (GUI). The user has direct control of parameters to control the rate of active neurons, their firing rate properties, and the duration of the recordings. The cell models are shipped with the software and recordings can be simulated on a single electrodes or a tetrode. It is relatively fast, but the cell model simulations (using `NEURON` [8]) are re-simulated for every recording.

Hagen et al. developed `ViSAPy` [19], a Python-based simulator that uses multi-compartment simulation of single neurons to generate spikes, network modeling of point-neurons in NEST [10] to generate synaptic inputs onto the spiking neurons, and experimentally fitted noise. `ViSAPy` runs a full network simulation in `NEURON` [8] and computes the extracellular potentials using `LFPy` [30, 18]. `ViSAPy` implements a Python application programming interface (API) which allows the user to set multiple parameters for the network simulation providing the synaptic input, the probe design, and the noise model generator. Cell models can be freely chosen and loaded using the `LFPy` package. Further, 1-dimensional drift can be incorporated in the simulations by shifting the electrodes over time [13]. Learning to use the software and, in particular, tailoring the specific properties of the resulting spike trains, for example burstiness, requires some effort by the user. As the running of `NEURON` simulations with biophysically detailed neurons can be computationally expensive, the use of `ViSAPy` to generate long-duration spike-sorting benchmarking data is boosted by access to powerful computers.

Mondragon et al. developed a Neural Benchmark Simulator (`NBS`) [35] extending the `NeuroCube` software. `NBS` extends the capability of `NeuroCube` for using user-specific probes, and it combines the spiking activity signals (from `NeuroCube`), with low-frequency activity signals, and artifacts libraries shipped with the code. The user can set different weight parameters to assemble the spiking, low-frequency, and artifact signals, but these three signal types are not modifiable.

Despite the existence of such tools for generating benchmarking data, their use in spike sorting literature has until now been limited, and the benchmarking and validation of spike sorting algorithms non-standardized and unsystematic. A natural question to ask is thus how to best stimulate the use of such benchmarking tools in the spike sorting community.

From a spike sorting developer perspective, we argue that an ideal extracellular simulator should be *i)* fast, *ii)* controllable, *iii)* biophysically detailed, and *iv)* easy to use. A fast simulator would enable spike sorter developers to generate a large and varied set of recordings to test their algorithms against and to improve their spike sorting methods. Controllability refers to the possibility to have direct control of features of the simulated recordings. The ideal extracellular spike simulator should include the possibility to use different cell models and types, to decide the firing properties of the neurons, to control the rate of spatio-temporal spike collisions, to generate recordings on different probe models, and to have full reproducibility of the simulated recordings. A biophysically detailed simulator should be capable of reproducing key physiological aspects of the recordings, including, but not limited to, bursting spikes, drifts between the electrodes and the neurons, and realistic noise profiles. Finally, to maximize the ease of use, the ideal extracellular simulator should be designed as an accessible and easy to learn software package. Preferably, the tool should be implemented with a graphical user interface (GUI), a command line interface (CLI), or with a simple application programming interface (API).

With these principles in mind, we present here `MEArec`, an open-source Python-based simulator. `MEArec` provides a fast, highly controllable, biophysically detailed, and easy to use framework to generate simulated extracellular recordings. In addition to producing benchmark datasets, we developed `MEArec` as a powerful tool that can serve as a testbench for optimizing existing and novel spike sorting methods. To facilitate this goal, `MEArec` allows users to explore how several aspects of recordings affect spike sorting, with full control of challenging features such as bursting activity, drifting, spatio-temporal synchrony, and noise effects, so that spike sorter developers can use it to help their algorithm design.

The source code for `MEArec` is on Github (https://github.com/alejoe91/MEArec) and the Python package is on `PyPi` (https://pypi.org/project/MEArec/). An extensive documentation is available (https://mearec.readthedocs.io/), and the code is tested with a continuous integration platform (https://travis-ci.org/). Moreover, all the datsets generated for this article and used to make figures are available on Zenodo (https://doi.org/10.5281/zenodo.3696926).

The article is organized as follows: in Section 2 we introduce the principles of `MEArec` and we show how to run simulations with the CLI and Python API. In Section 3 we explain the different features available in `MEArec`, including the capability of simulating recordings for MEAs, reproducing bursting behavior, controlling spatio-temporal overlaps, reproducing drifts, and replicating biological noise characteristics. In Section 4 we present the use of MEArec as a testbench for spike sorting development, and its integration with the SpikeInterface framework [4]. In Section 5 we document the simulation outputs and how to save and load them with the `MEArec` API. Finally, in Section 6 we discuss the presented software and contextualize it with respect to the state of the art.

## 2 Getting started with MEArec

We start by describing the principle of the `MEArec` simulator and showing examples on how to get started with the simulations.

The simulation is split in two phases: *templates generation* (Figure 1A) and *recordings generation* (Figure 1B). Templates (or extracellular action potentials - EAPs) are generated using biophysically realistic cell models which are positioned in the surroundings of a probe model. The templates generation phase is further divided into an *intracellular* and an *extracellular* simulation. During the intracellular simulation, each cell model is stimulated with a constant current and transmembrane currents of action potentials are computed (using `NEURON` [8]) and stored to disk (the *intracellular* simulation is the most time consuming part and storing its output to disk enables one to run it only once). The extracellular simulation uses the `LFPy` package [30, 18] to compute extracellular potentials generated at the electrodes’ locations using the well-established line-source approximation (see Supplementary Methods – *Templates generation* –for details). In particular, the cell morphology is loaded and shifted to a random position around the probe. Additionally, the user can add different rotations to the models. When the cell model is shifted and rotated, the previously computed and stored transmembrane currents are loaded and the EAP is computed. This step is repeated several times for each cell model, for different positions and rotations. The templates generation phase outputs a library of a large variety of extracellular templates, which can then be used to build the recordings. The templates generation phase is the most time consuming, but the same templates library can be used to generate multiple recordings. It is therefore recommended to simulate many more templates than needed by a single recording, so that the same template library can be used to simulate a virtually infinite number of recordings.

**Figure 1:**
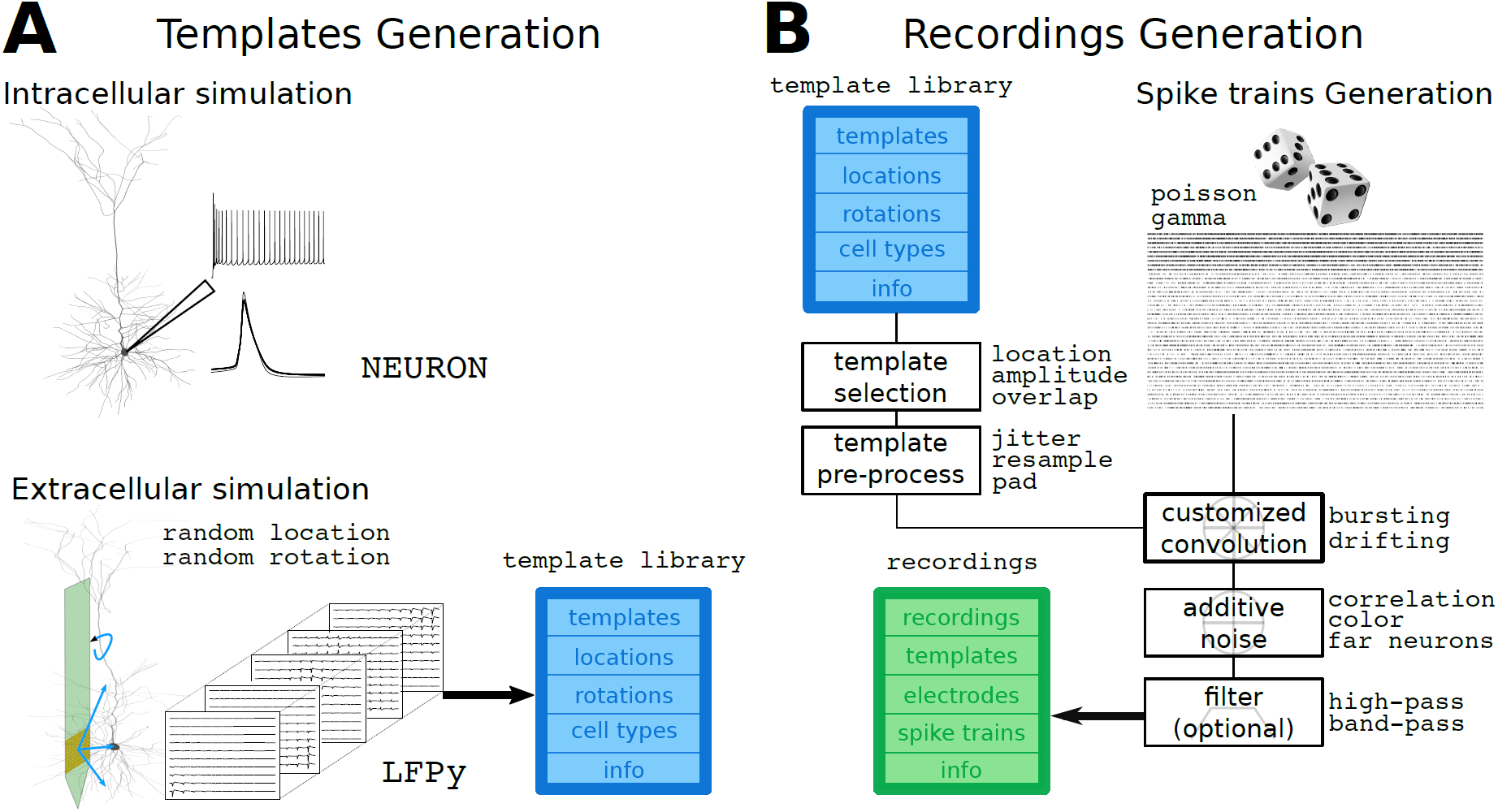
Overview of the `MEArec` software. The simulation is divided in two phases: *templates generation* and *recordings generation*. (A) The *templates generation* phase is split in an intracellular and extracellular simulation. The intracellular simulation computes, for each available cell model, the transmembrane currents generated by several action potentials. In the extracellular simulation, each cell model is randomly moved and rotated several times and the stored currents are loaded to the model to compute the extracellular action potential, building a *template library*. (B) The *recordings generation* phase combines templates selected from the *template library* and randomly generated spike trains. Selected templates are pre-processed before a customized convolution with the spike trains. Additive noise is added to the output of the convolution, and the recordings can be optionally filtered.

`MEArec`, at installation, comes with 13 layer 5 cortical cell models from the Neocortical Microcircuit Portal [42]. This enables the user to dive into simulations without the need to download and compile cell models. On the other hand, the initial set of cell models can be easily extended as outlined in the Supplementary Methods – *Templates generation*.

To generate 30 extracellular spikes (also referred as templates) per cell model recorded on a shank tetrode probe, the user can simply run this command:

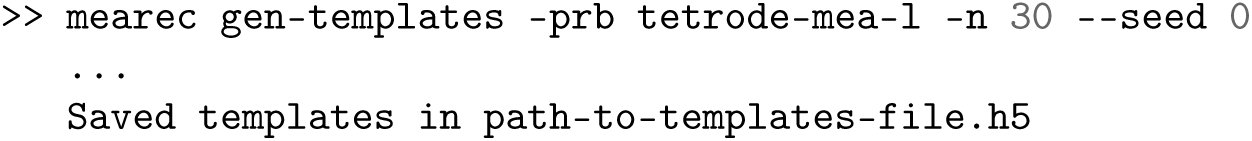

The `-prb` option allows for choosing the probe model, `-n` controls the number of templates per cell model to generate, and the `--seed` option is used to ensure reproducibility and if it is not provided, a random seed is chosen. In both cases, the seed is saved in the `HDF5` file, so that the same templates can be replicated.

Recordings are then generated by combining templates selected with user-defined rules (based on minimum distance between neurons, amplitudes, spatial overlaps, and cell-types) and by simulating spike trains (Supplementary Methods – *Recordings generation* – for details on spike trains generation and template selection). Selected templates and spike trains are assembled using a customized (or modulated) convolution, which can replicate interesting features of spiking activity such as bursting and drift. After convolution, additive noise is generated and added to the recordings. Finally, the output recordings can be optionally filtered with a band-pass or a high-pass filter. Note that filtering the recordings will affect the shape and amplitude of the spike waveforms, but this is a common procedure in spike sorting to remove lower frequency components.

Recordings can be generated with the CLI as follows:

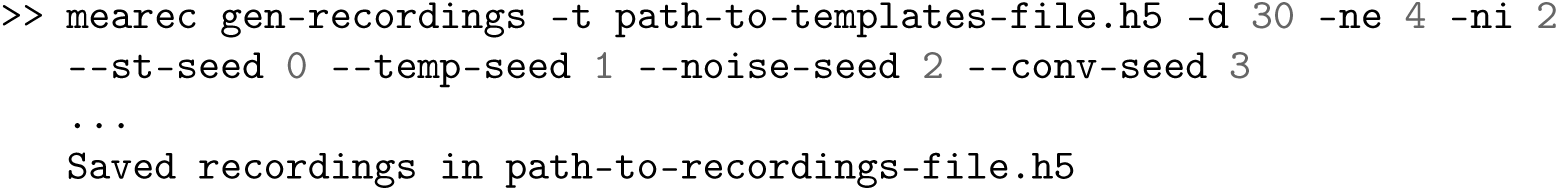

The `gen-recordings` command combines the selected templates from 4 excitatory cells (`-ne 4`) and 2 inhibitory cells (`-ni 2`), that usually have a more narrow spike waveform and a higher firing rate, with randomly generated spike trains. The duration of the output recordings is 30 seconds (`-d 30`). In this case, four random seeds control the spike train random generation (`--st-seed 0`), the template selection (`--temp-seed 1`), the noise generation (`--noise-seed 2`), and the convolution process (`--conv-seed 3`). Figure 2 shows one second of the generated recordings (A), the extracted waveforms and the mean waveforms for each unit on the electrode with the largest peak (B), and the principal component analysis (PCA) projections of the waveforms on the tetrode channels.

**Figure 2:**
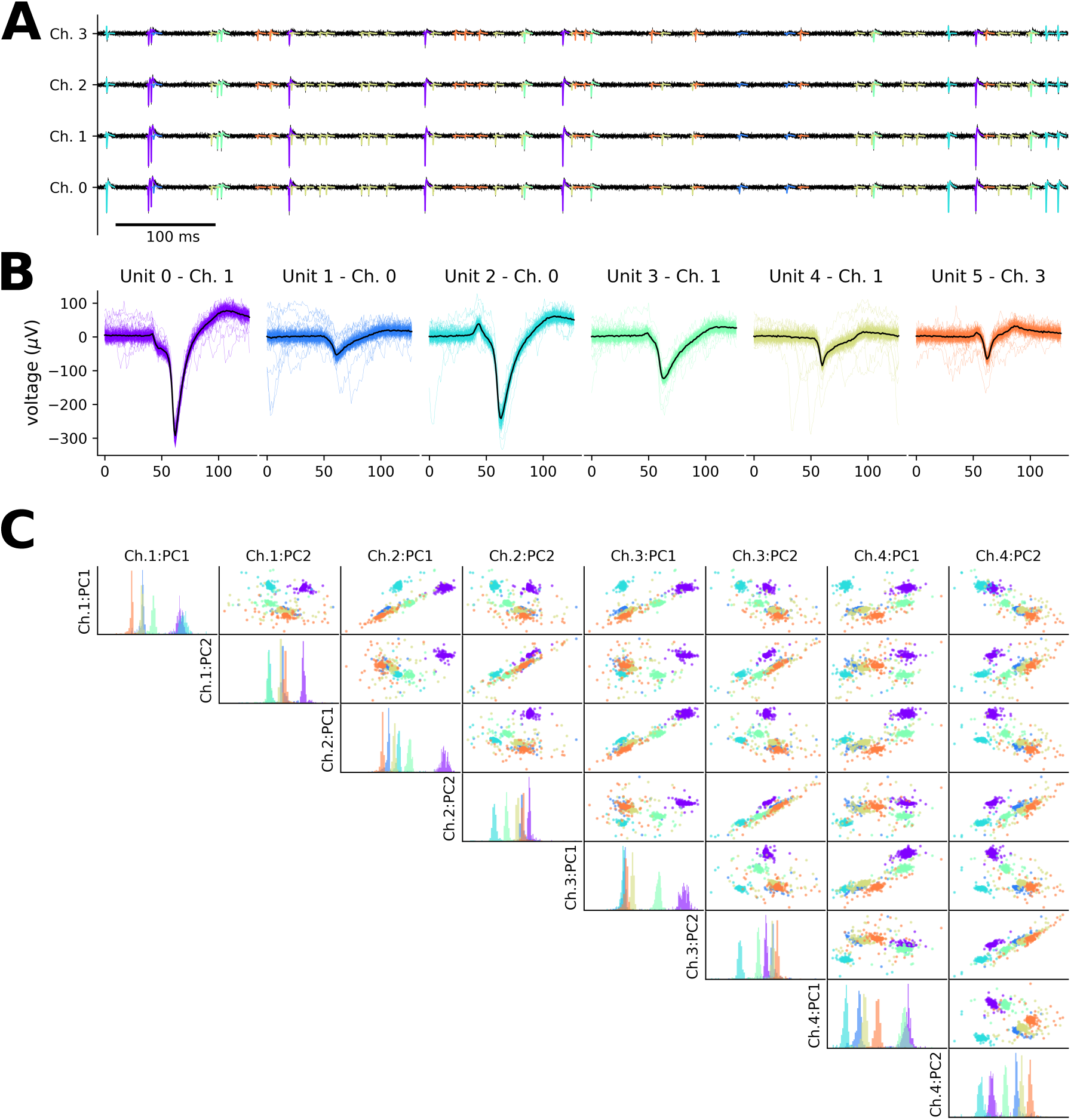
Example of simulated tetrode recording. (A) One second of the recording timeseries on the four tetrode channels. The templates for the different units are overlapped to the recording traces in different colors. (B) Extracted waveforms on the channel with the largest amplitude for the six units in the recordings. (C) PCA projections on the first two PC components of the four tetrode channels. Each color corresponds to a neuron. The diagonal plots display the histograms of the PC projection on the corresponding channel.

`MEArec` also implements a convenient Python API, which is run internally by the CLI commands. For example, the following snippet of code implements the same commands shown above for generating templates and recordings:

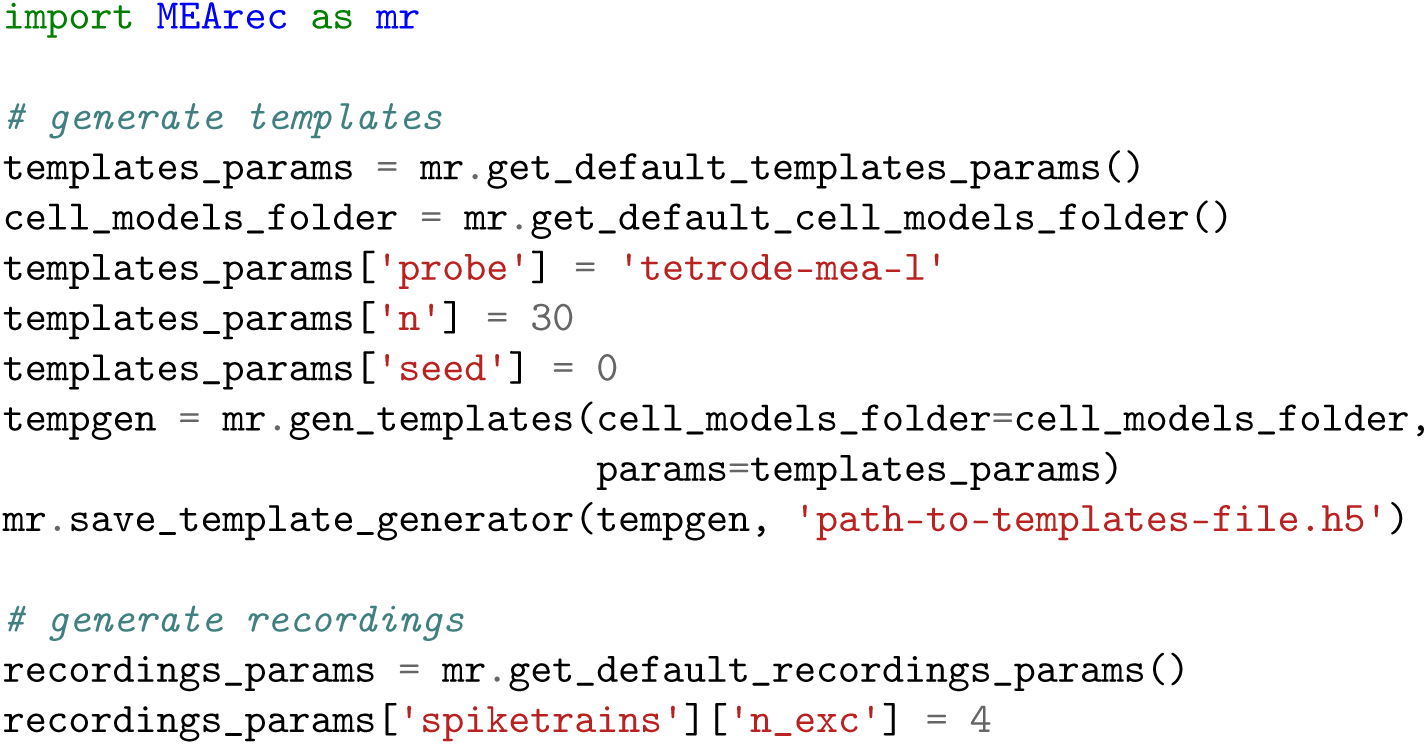
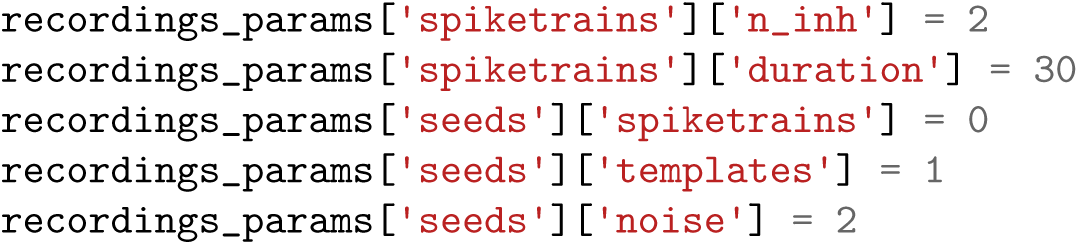

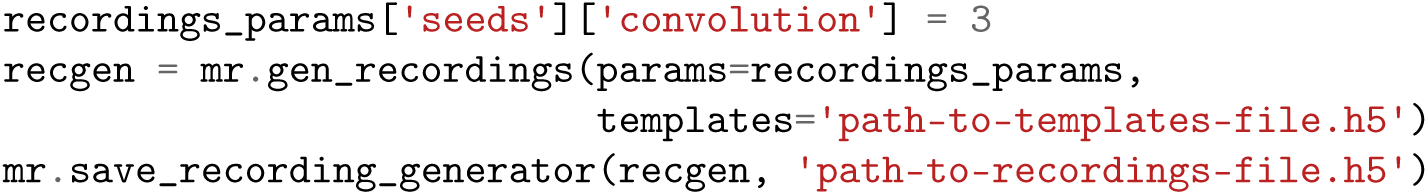

Moreover, the Python API implements plotting functions to visually inspect the simulated templates and recordings. For example, Figure 2 panels were generated using the `plot_recordings()` (A), `plot_waveforms()` (B), and `plot_pca_map()` (C) functions.

`MEArec` is designed to allow for full customization, transparency, and reproducibility of the simulated recordings. Parameters for the templates and recordings generation are accessible by the user and documented, so that different aspects of the simulated signals can be finely tuned (see Supplementary Methods for a list of parameters and their explanation). Moreover, the implemented command line interface (CLI) and simple Python API, enables the user to easily modify parameters, customize, and run simulations.

Finally, `MEArec` permits to manually set several random seeds used by the simulator to make recordings fully reproducible. This feature also enables one to study how separate characteristics of the recordings affect the spike sorting performance. As an example, we will show in the next sections how to simulate a recording sharing all parameters, hence with exactly the same spiking activity, but with different noise levels or drifting velocities.

## 3 MEArec features

### 3.1 Generation of realistic Multi-Electrode Array recordings

The recent development of Multi-Electrode Arrays (MEAs) enables researchers to record extracellular activity at very high spatio-temporal density both for *in vitro* [2, 14] and *in vivo* applications [37, 27]. The large number of electrodes and their high density can result in challenges for spike sorting algorithms. It is therefore important to be able to simulate recordings from these kind of neural probes.

To deal with different probe designs, `MEArec` uses another Python package (`MEAutility` - https://meautility.readthedocs.io/), that allows users to easily import several available probe models and to define custom probe designs. Among others, `MEAutility` include Neuropixels probes [27], Neuronexus commercial probes (http://neuronexus.com/products/neural-probes/), and a wide variety of square MEA designs with different contact densities (the list of available probes can be found using the `mearec available-probes` command).

Similarly to the tetrode example, we first have to generate templates for the probes. These are the commands to generate templates and recordings for a Neuropixels design with 128 electrodes (`Neuropixels-128`). The recordings contain 60 neurons, 48 excitatory and 12 inhibitory. With similar commands, we generated templates and recordings for a Neuronexus probe with 32 channels (A1×32-Poly3-5mm-25s-177-CM32 - `Neuronexus-32`) with 20 cells (16 excitatory and 4 inhibitory), and a square 10×10 MEA with 15 µm inter-electrode-distance (`SqMEA-10-15`) and 50 cells (40 excitatory and 10 inhibitory).

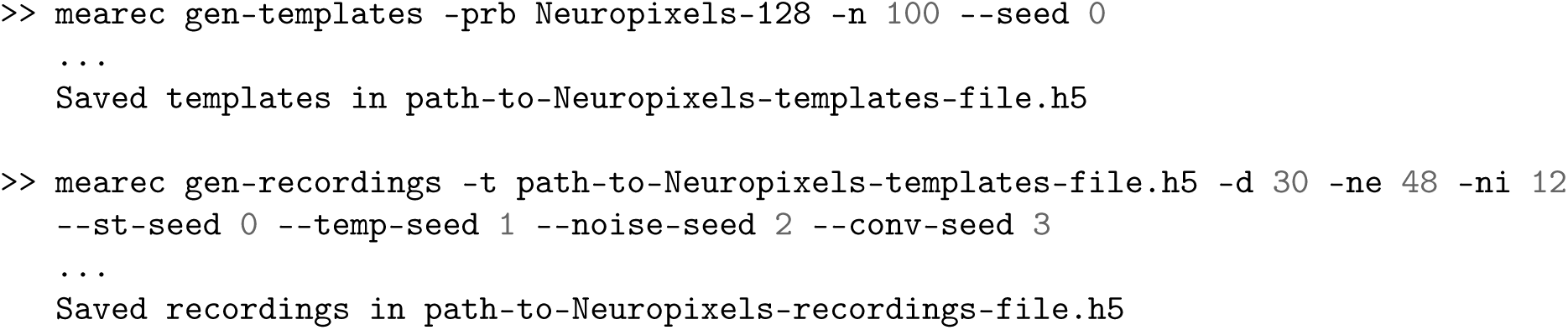

Figure 3 shows the three above-mentioned probes (A), a sample template for each probe design (B), and one-second snippets of the three recordings (C-D-E), with zoomed in windows to highlight spiking activity.

**Figure 3:**
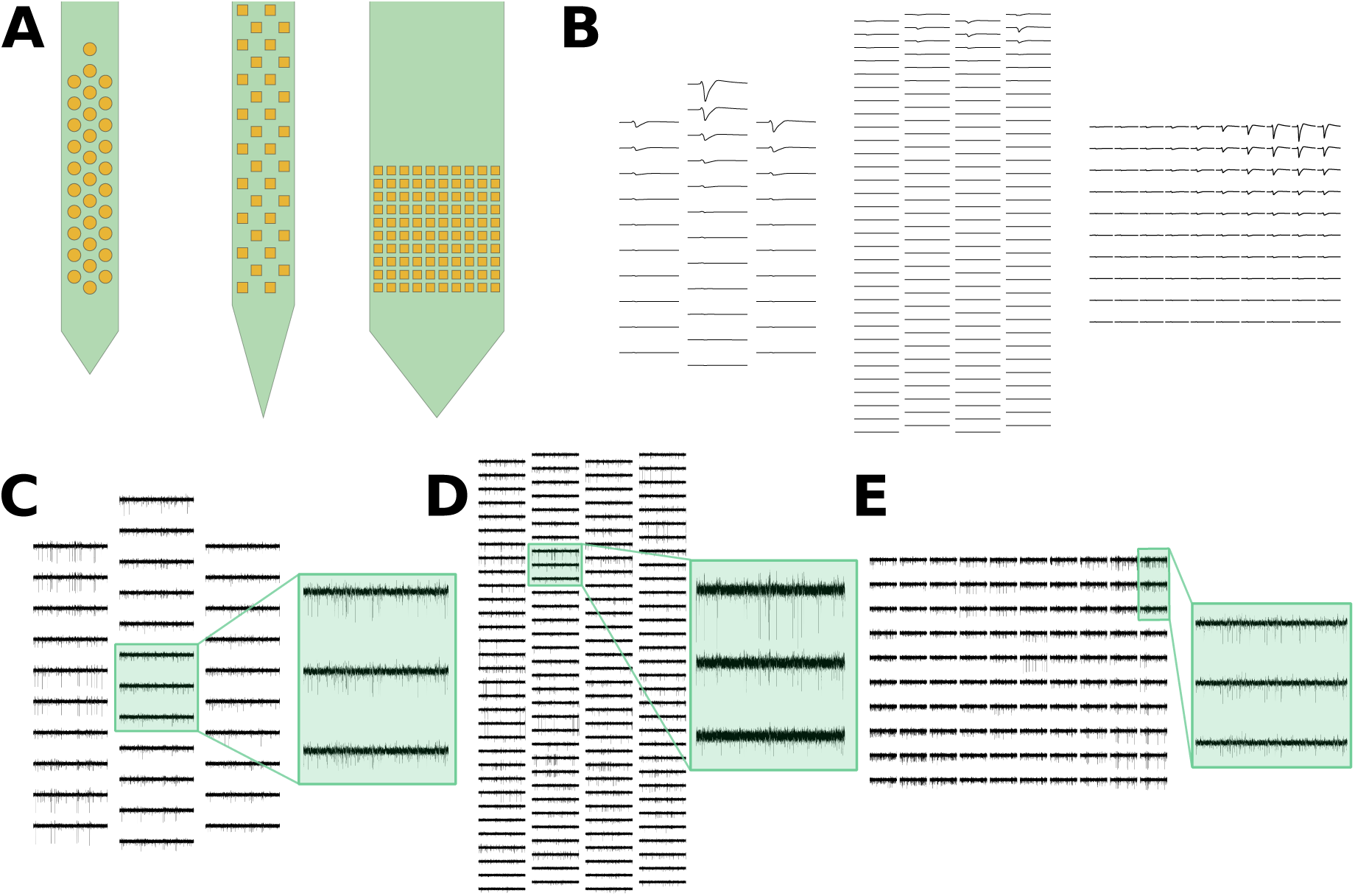
Generation of high-density multi-electrode array recordings. (A) Example of three available probes: a commercial Neuronexus probe (left), the Neuropixels probe (middle), and a high-density square MEA. (B) Sample templates for each probe design. (C-D-E) One-second snippets of recordings from the Neuronexus probe (C), the Neuropixels probe (D), and the square MEA probe (E). The highlighted windows display the activity over three adjacent channels and show how the same spikes are seen on multiple sites.

While all the recordings shown so far have been simulated with default parameters, several aspects of the spiking activity are critical for spike sorting. In the next sections, we will show how these features, including bursting, spatio-temporal overlapping spikes, drift, and noise assumptions can be explored with `MEArec` simulations.

### 3.2 Bursting modulation of spike amplitude and shape

Bursting activity is one of the most complicated features of spiking activity that can compromise the performance of spike sorting algorithms. When a neuron bursts, i.e., fires a rapid train of action potentials with very short inter-spike intervals, the dynamics underlying the generation of the spikes changes over the bursting period [21]. While the bursting mechanism has been largely studied with patch-clamp experiments, combined extracellular-juxtacellular recordings [1] and computational studies [19] suggest that during bursting, extracellular spikes become lower in amplitude and wider in shape.

In order to simulate this property of the extracellular waveforms in a fast and efficient manner, templates can be modulated both in amplitude and shape during the convolution operation, depending on the spiking history.

To demonstrate how bursting is mimicked, we built a toy example with a constant spike train with 10 ms inter-spike-interval (Figure 4A). A *modulation value* is computed for each spike and it is used to modulate the waveform for that event by scaling its amplitude, and optionally stretching its shape. The blue dots show the default modulation (bursting disabled), in which the modulation values are drawn from a Gaussian distribution with unitary mean to add some physiological variation to the spike waveforms. When bursting is enabled (by setting the `bursting` parameter to true), the modulation values are computed based on the spike history, and it depends on the number of consecutive spikes in a bursting event and their average inter-spike-intervals (see Supplementary Methods – *Recordings generation - Modulated convolution* – for details on the modulation values calculation).

**Figure 4:**
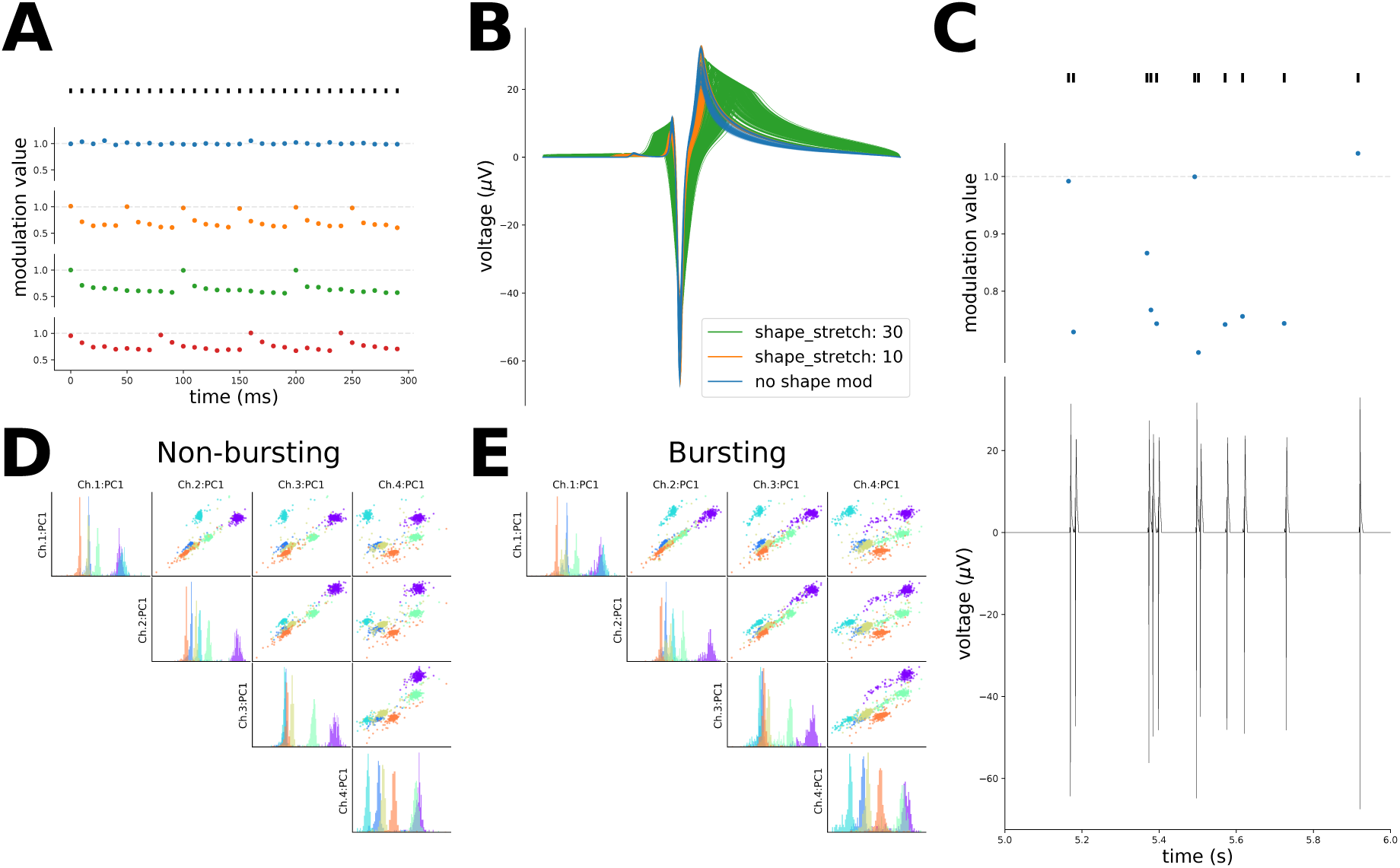
Bursting behavior. (A) Modulation values computation for a sample spike train of 300 ms with constant inter-spike-intervals of 10 ms. The blue dots show the modulation values for each spike when bursting is not activated: each value is drawn from a *𝒩* (1, 0.05^2^) distribution. When bursting is activated, a bursting event can be limited by the maximum number of spikes (orange - 5 spikes, green - 10 spikes), or by the maximum bursting event duration (red - 75 ms). (B) Modulated templates. The blue lines show templates modulated in amplitude only. The orange and green lines display the same templates with added shape modulation. (C) Modulation in tetrode recordings. The top panel shows spikes in a one-second period. The middle panel displays the modulation values for those spikes. The bottom panel shows the modulated template on the electrode with the largest peak after convolution. (D-E) PCA projections on the first principal component for the tetrode recordings without bursting (the same as Section 2) (D) and re-simulated with bursting (E) and shape modulation enabled. Note that the PCA projections were computed in both cases from the waveforms without bursting. The clusters, with bursting, become more spread and harder to separate than without bursting.

Bursting events can be either controlled by the maximum number of spikes making a burst (orange dots - 5 spikes per burst; green dots - 10 spikes per burst) or by setting a maximum bursting duration (red dots - maximum 75 ms). Note that in Figure 4A the spike train is constant just to illustrate the computation of the modulation values. In actual simulations, instead, the modulation values will depend on the firing rate and the timing between spikes.

By default, spikes are only modulated in amplitude. The user can also enable shape modulation by setting the `shape_mod` parameter to true. The modulation value, computed for each spike, controls both the amplitude scaling and shape modulation of the spike event. For amplitude modulation, the amplitude of the spike is simply multiplied by the modulation value. Additionally, when shape modulation is enabled, the waveform of each spike is also stretched. The `shape_stretch` parameter controls the overall amount of stretch, but that actual stretch of single waveforms depends on the modulation value computed for each spike. In Figure 4B, examples of bursting templates are shown. The blue traces display templates only modulated in amplitude, i.e., the amplitude is scaled by the modulation value. The orange and green traces, instead, also present shape modulation, with different values of the `shape_stretch` parameter (the higher the `shape_stretch`, the more stretched waveforms will be). We refer to the Supplementary Methods – *Recordings generation - Modulated convolution* – for further details on amplitude and shape modulation.

Figure 4C shows a one-second snippet of the tetrode recording shown previously after bursting modulation is activated. The top panel shows the spike events, the middle one displays the modulation values computed for each spike, and the bottom panel shows the output of the modulated convolution between one of the templates (on the electrode with the largest amplitude) and the spike train.

Figure 4D and Figure 4E show the waveform projections on the first principal component of each channel for the tetrode recording shown in Section 2 with and without bursting enabled, respectively. In this case all neurons are bursting units and this causes a stretch in the PCA space, which is a clear complication for spike sorting algorithms. Note that shape modulation does not affect all neurons by the same amount, since it depends on the spike history and therefore on the firing rate.

### 3.3 Controlling spatio-temporal overlaps

Another complicated aspect of extracellular spiking activity that can influence spike sorting performance is the occurrence of overlapping spikes. While temporal overlapping of events on spatially separated locations can be solved with feature masking [44], spatio-temporal overlapping can cause a distortion of the detected waveform, due to the superposition of separate spikes. Some spike sorting approaches, based on template-matching, are designed to tackle this problem [40, 47, 11].

In order to evaluate to what extent spatio-temporal overlap affects spike sorting, `MEArec` allows the user to set the number of spatially overlapping templates and to modify the synchrony rate of their spike trains. In Figure 5 we show an example of this on a `Neuronexus-32` probe (see Figure 3A). The recording was constructed with two excitatory and spatially overlapping neurons, whose templates are shown in Figure 5A (see Supplementary Methods – *Recordings generation - Overlapping spikes and spatio-temporal synchrony* – for details on the spatial overlap definition). The spike synchrony rate can be controlled with the `sync_rate` parameter. If this parameter is not set (Figure 5B - left), some spatio-temporal overlapping spikes are present (red events). If the synchrony rate is set to 0, those spikes are removed from the spike trains (Figure 5B - middle). If set to 0.05, i.e., 5% of the spikes will be spatio-temporal collisions, events are added to the spike trains to reach the specified synchrony rate value of spatio-temporal overlap. As shown in Figure 5C, the occurrence of spatio-temporal overlapping events affects the recorded extracellular waveform: the waveforms of the neurons, in fact, get summed and might be mistaken for a separate unit by spike sorting algorithms when the spikes are overlapping.

**Figure 5:**
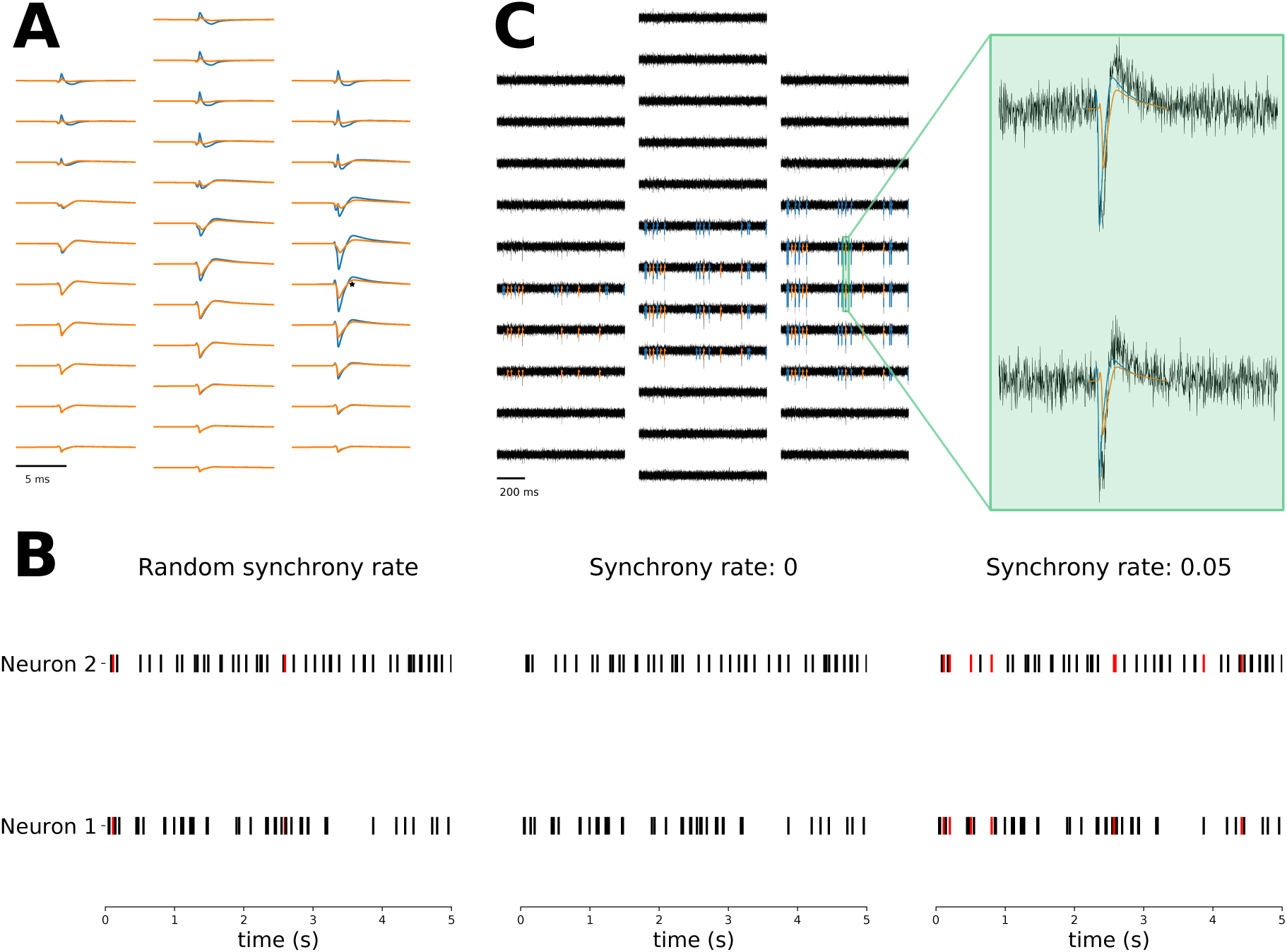
Controlling spatio-temporal overlapping spikes. (A) Example of two spatially overlapping templates. The two templates are spatially overlapping because on the electrode with the largest signal (depicted as an black asterisk) for the blue template, the orange template has an amplitude greater than the 90% of its largest amplitude. (B) Without setting the synchrony rate, the random spike trains (left) present a few spatio-temporal collisions (red events). When setting the synchrony rate to 0 (middle), the spatio-temporal overlaps are removed. When the synchony rate is set to 0.05 (right), spatio-temporal overlapping spikes are added to the spike trains. (C) One-second snippet of the recording with 0.05 synchrony. In the magnified window, a spatio-temporal overlapping event is shown: the collision results in a distortion of the waveform.

The possibility of reproducing and controlling this feature of extracellular recordings within `MEArec` could aid in the development of spike sorters which are robust to spatio-temporal collisions.

### 3.4 Generating drifting recordings

When extracellular probes are inserted in the brain, especially for acute experiments, the neural tissue might slowly move with respect to the electrodes. This phenomenon is known as drift. Drift can be due to a slow relaxation of the tissue (slow drift) or to fast re-adjustments of the tissue, for example due to an abrupt motion of the tissue (fast drift). These two types of drifts can also be observed in tandem [39].

Drifting units are particularly critical for spike sorting, as the waveform shapes change over time due to the relative movement between the neurons and the probe. New spike sorting algorithms have been developed to specifically tackle the drifting problem (e.g. `Kilosort2` [39], `IronClust` [26]).

In order to simulate drift in the recordings, we first need to generate drifting templates:

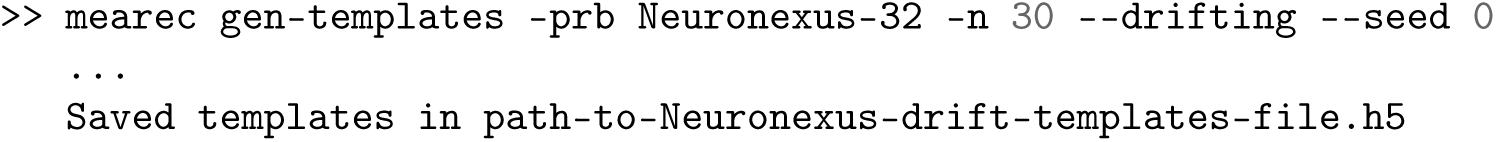

Drifting templates are generated by choosing an initial and final soma position with user-defined rules (see Supplementary Methods – *Template generation - Drifing templates* – for details) and by moving the cell along the line connecting the two positions for a defined number of constant drifting steps that span the segment connecting the initial and final positions (30 steps by default). An example of a drifting template is depicted in Figure 6A, alongside with the drifting neuron’s soma locations for the different drifting steps.

**Figure 6:**
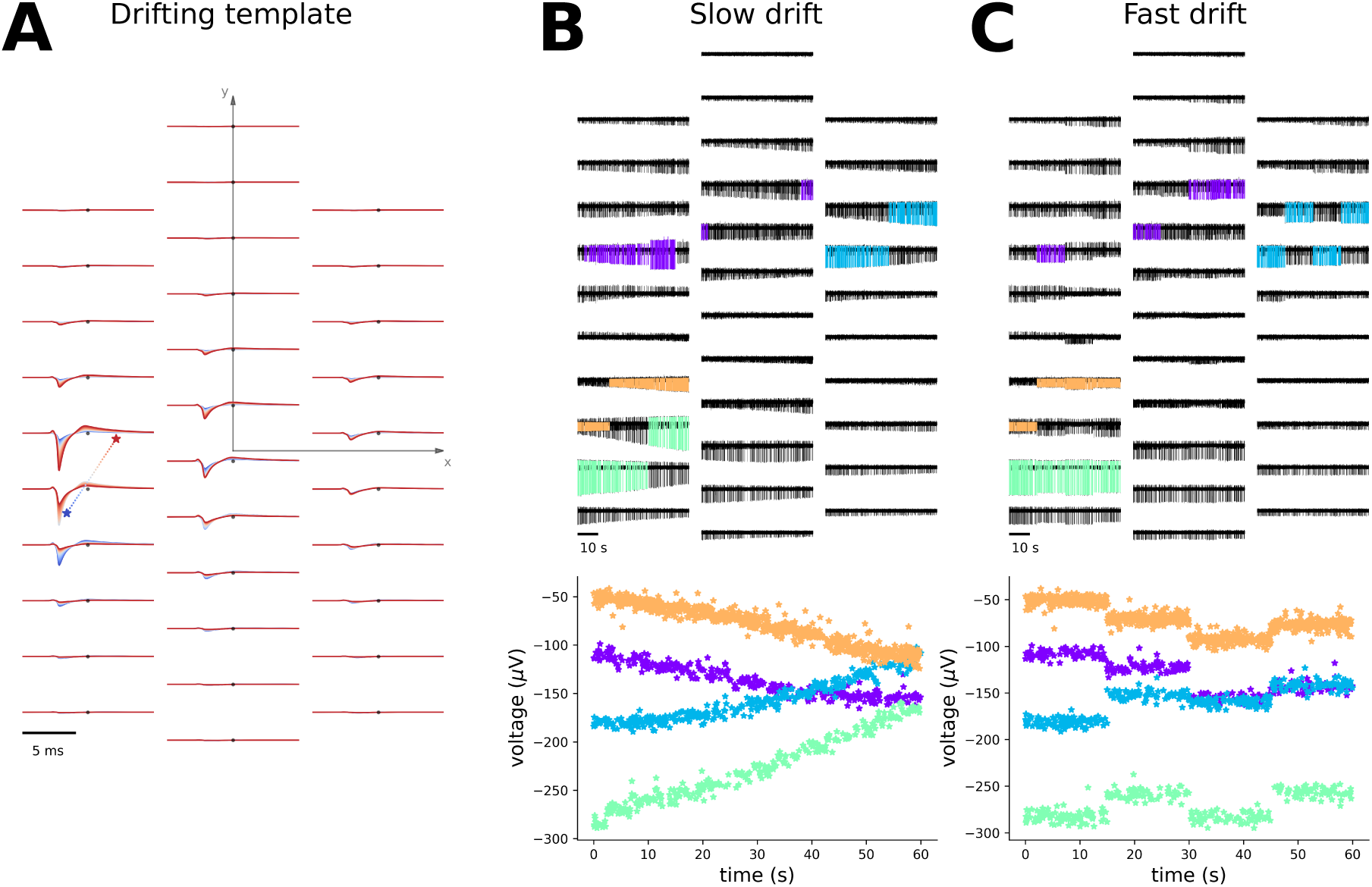
Drifting. (A) Example of a drifting template. The colored asterisks on the left show the trajectory from the initial (blue large asterisk) to the final (red large asterisk) neuron positions. The positions are in the x-y coordinates of the probe plane, and the electrode locations are depicted as black dots. The corresponding templates are displayed at the electrode locations with the same colormap, showing that the template peak is shifted upwards following the soma position. (B) Slow drift. (top) 60-second slow drifting recording with four neurons moving at a velocity of 20 µm*/min*. Templates on the largest electrode are superimposed in different colors on the recordings. Note that the maximum channel changes over time. (bottom) Amplitude of the waveforms over time on the electrode with the largest initial peak. (C) Fast drift. (top) 60-second fast drifting recording with four neurons undergoing a fast drift event every 15 s. Templates on the largest electrode are superimposed in different colors on the recordings. Also for fast drifts, the maximum channel changes over time. (bottom) Amplitude of the waveforms over time on the electrode with the largest initial peak.

Once a library of drifting templates is generated, drifting recordings can be simulated. `MEArec` allows users to simulate recordings with three types of drift modes: *slow, fast*, and *slow+fast*. When slow drift is selected, the drifting template is selected over time depending on the initial position and the drifting velocity (5 µm*/min* by default). If the final drifting position is reached, the drift direction is reversed. For fast drifts, the position of a drifting neuron is shifted abruptly with a user-defined period (every 20 s by default). The new position is chosen so that the difference in waveform amplitude of the drifting neuron on its current maximum channel remains within user-defined limits (5-20 µV by default), in order to prevent from moving the neuron too far from its previous position. The slow+fast mode combines the slow and fast mechanisms.

In Figure 6B and C we show examples of slow drift and fast drift, respectively. In the top panel the recordings are displayed, with superimposed drifting templates on the electrode with the largest peak. Note that the maximum channel can change over time due to drift. In the bottom panels, instead, the amplitude of the waveforms on the channels with the initial largest peak for each neuron are shown over time. Slow drift causes the amplitude to slowly vary, while for fast drifts we observe more abrupt changes when a fast drift event occurs. In the *slow+fast* drift mode, these two effects are combined.

### 3.5 Modeling experimental noise

Spike sorting performance can be greatly affected by noise in the recordings. Many algorithms first use a spike detection step to identify putative spikes. The threshold for spike detection is usually set depending on the noise standard deviation or median absolute deviation [41]. Clearly, recordings with larger noise levels will result in higher spike detection thresholds, hence making it harder to robustly detect lower amplitude spiking activity. In addition to the noise amplitude, other noise features can affect spike sorting performance: some clustering algorithms, for example, assume that clusters have Gaussian shape, due to the assumption of an additive normal noise to the recordings. Moreover, the noise generated by biological sources can produce spatial correlations in the noise profiles among different channels and it can be modulated in frequency [7, 43].

To investigate how the above-mentioned assumptions on noise can affect spike sorting performance, `MEArec` can generate recordings with several noise models. Figure 7 shows 5-second spiking-free recordings of a tetrode probe for five different noise profiles that can be generated (A - recordings, B - spectrum, C - channel covariance, D - amplitude distribution).

**Figure 7:**
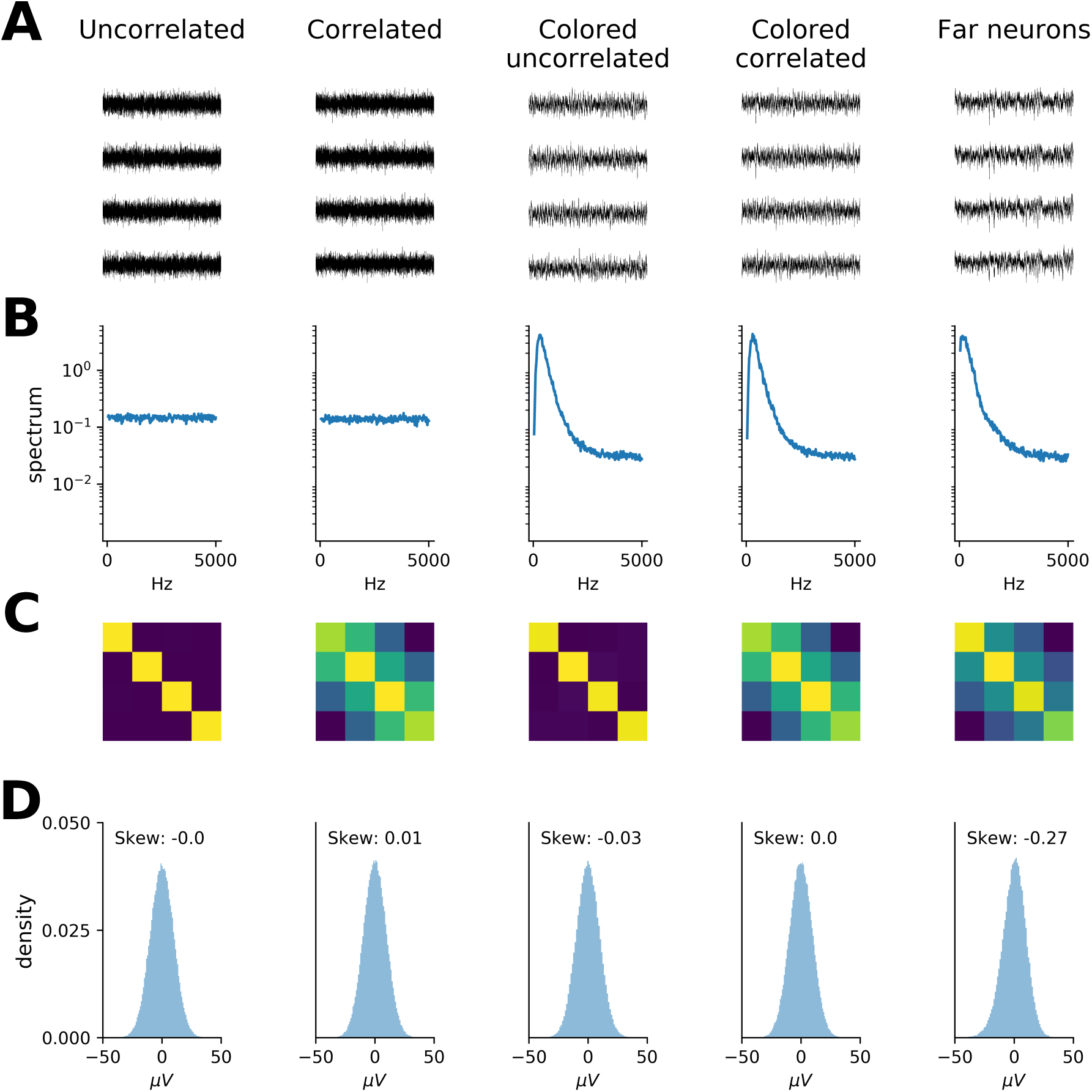
Noise models. The 5 columns refer to different noise models: 1) Uncorrelated Gaussian noise, Distance-correlated Gaussian noise, 3) Colored uncorrelated Gaussian noise, 4) Colored distance-correlated Gaussian noise, and 5) Noise generated by distant neurons. (A) One-second spiking-free recording. (B) Spectrum of the first recording channel between 10 and 5000 Hz. (C) Covariance matrix of the recordings. (D) Distribution of noise amplitudes for the first recording channel. The different noise models vary in the spectrum, channel correlations, and amplitude distributions.

The first column shows uncorrelated Gaussian noise, which presents a flat spectrum, a diagonal covariance matrix, and a symmetrical noise amplitude distribution. In the recording in the second column, spatially correlated noise was generated as a multivariate Gaussian noise with a covariance matrix depending on the channel distance. Also in this case, the spectrum (B) presents a flat profile and the amplitude distribution is symmetrical (D), but the covariance matrix shows a correlation depending on the inter-electrode distance. As previous studies showed [7, 43], the frequency content of extracellular noise is not flat, but its spectrum is affected by the spiking activity of distant neurons, which appear in the recordings as below-threshold *biological* noise. To reproduce the spectrum profile that is observed in experimental data, `MEArec` allows coloring the noise spectrum of Gaussian noise with a second order infinite impulse response (IIR) filter (see Supplementary Methods – *Recordings generation - Noise models and post-processing* – for details). Colored noise represents an efficient way of obtaining the desired spectrum, as shown in the third and fourth columns of Figure 7, panel B. Distance correlation is maintained (panel C - fourth column), and the distribution of the noise amplitudes is symmetrical. Finally, a last noise model enables one to generate activity of distant neurons. In this case, noise is built as the convolution between many neurons (300 by default) whose template amplitudes are below an amplitude threshold (10 µV by default). A Gaussian noise floor is then added to the resulting noise, which is scaled to match the user-defined noise level. The *far-neurons* noise profile is shown in the last column of Figure 7. While the spectrum and spatial correlation of this noise profile are similar to the ones generated with a colored, distance-correlated noise (4th column), the shape of the noise distribution is skewed towards negative values (panel D), mainly due to the negative contribution of the action potentials.

The capability of `MEArec` to simulate several noise models enables spike sorter developers to assess how different noise profiles affect their algorithms and to modify their methods to be insensitive to specific noise assumptions.

## 4 Testbench for spike sorting development and evaluation

In the previous sections, we have shown several examples on how `MEArec` is capable of reproducing several aspects of extracellular recordings which are critical for spike sorting performance, in a fully reproducible way. The proposed design and its integration with a spike sorting evaluation framework called `SpikeInterface` [4] enables developers to actively include customized simulations in the spike sorting development phase.

Due to its speed and controllability, we see `MEArec` as a *testbench*, rather than a *benchmark* tool. We provide here a couple of examples. In Figure 8A, we show a one-second section of recordings simulated on a `Neuronexus-32` probe with fixed parameters and random seeds regarding template selection and spike train generation, but with four different levels of additive Gaussian noise, with standard deviations of 5, 10, 20, and 30 µV. The traces show the same underlying spiking activity, so the only variability in spike sorting performance will be due to the varying noise levels. Similarly, in Figure 8B, 1-minute drifting recordings were simulated with three different drifting velocities. The recordings show that for low drifting speeds the waveform changes are almost not visible (green traces), while for faster drifts (orange and blue traces), the waveform changes over time become more important.

**Figure 8:**
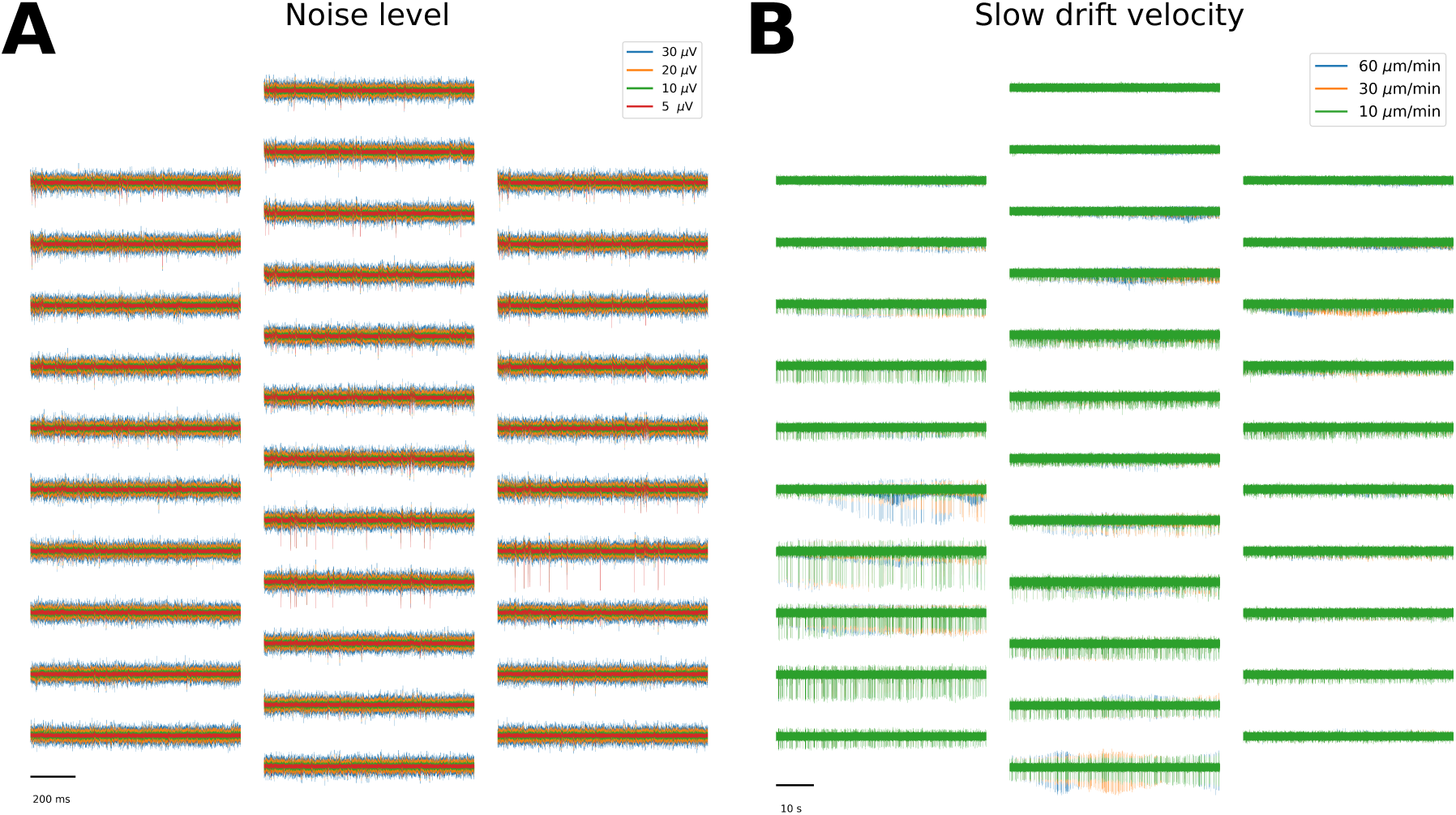
`MEArec` as testbench platform for spike sorting. (A) Four one-second snippet of recordings generated with a different noise level parameter (5 - red, 10 - green, 20 - orange, and 30 µV - blue). The underlying spiking activity is exactly the same for all recordings, and the only difference lies in the standard deviation of the underlying uncorrelated Gaussian noise. (B) Three slow drifting recordings generated with a different drifting velocity parameter (10 - green, 30 - orange, and 60 µm*/min* - blue). Also in this case, the underlying spiking activity is the same, but it can be observed how the different speeds result in a modification of waveforms over time.

The capability of `MEArec` of reproducing such behaviors in a highly controlled manner could aid in the design of specific tests for measuring and quantifying the ability of a spike sorting software to deal with specific complexities in extracellular recordings. Other examples include simulating a recording with increasing levels of bursting in order to measure to what extent bursting units are correctly clustered, or changing the synchrony rate of spatially overlapping units to assess how much spatio-temporal collisions affect performance.

### 4.1 Integration with SpikeInterface

We have recently developed `SpikeInterface` [4], a Python-based framework for running several spike sorting algorithms, comparing, and validating their results. `MEArec` can be easily interfaced to `SpikeInterface` so that simulated recordings can be loaded, spike sorted, and benchmarked with a few lines of code. In the following example, a `MEArec` recording is loaded, spike sorted with `Mountainsort4` [9] and `Kilosort2` [39], and benchmarked with respect to the ground-truth spike times available from the `MEArec` simulation:

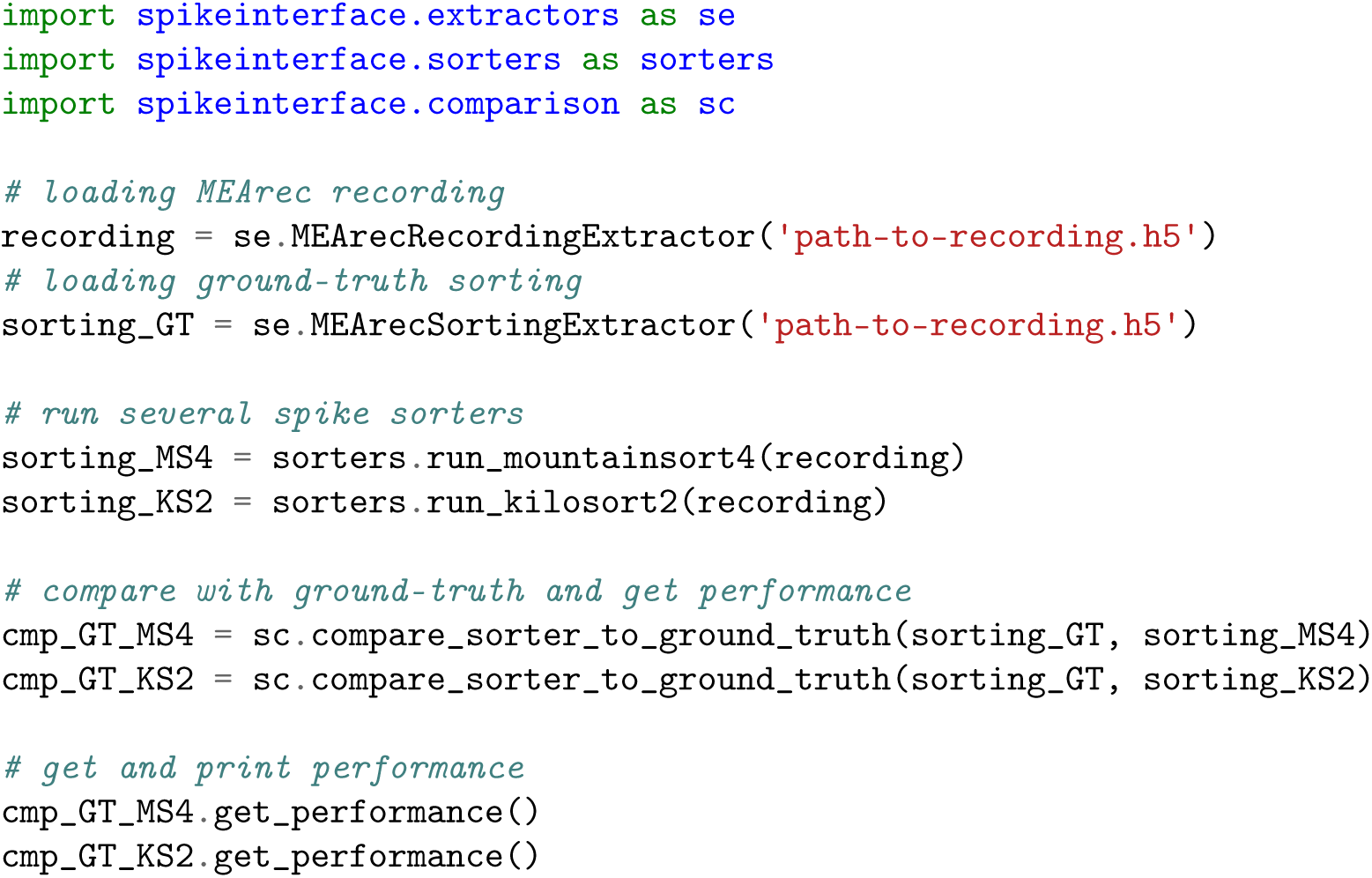

The `get_performance()` function returns the accuracy, precision, and recall for all the ground-truth units in the MEArec recording. For further details on these metrics and a more extensive characterization of the comparison we refer to the `SpikeInterface` documentation and article [4].

The combination of `MEArec` and `SpikeInterface` represents a powerful tool for systematically testing and comparing spike sorter performances with respect to several complications of extracellular recordings. `MEArec` simulations, in combination with `SpikeInterface`, are already being used by other groups to benchmark and compare spike sorting algorithms by the `SpikeForest` project [31].

### 4.2 Performance considerations

As a testbench tool, the speed requirement has been one of the main design principle of `MEArec`. In order to achieve high speed, most parts of the simulation process are fully parallelized. As shown in Figure 1, the simulations are split in templates and recordings generation. The templates generation phase is the most time consuming, but the same template library can be used to generate several recordings. This phase is further split in two sub-phases: the intracellular and extracellular simulations. The former only needs to be run once, as it generates a set of cell model-specific spikes that are stored and then used for extracellular simulations, which is instead probe specific.

We present here run times for the different phases of the templates generation and for the recordings generation. All simulations were run on an Ubuntu 18.04 Intel(R) Core(TM) i7-6600U CPU @ 2.60GHz, with 16 GB of RAM.

The intracellular simulation run time for the 13 cell models shipped with the software was ∼ 130 seconds (∼ 10 seconds per cell model).

Run times for extracellular simulations for several probe types, number of templates in the library, and drifting templates are shown in the *Templates generation* section of Table 1. The run times for this phase mainly depend on the number of templates to be generated (N templates column), on the minimum amplitude of accepted templates (Min. amplitude column, see Supplementary Methods – *Templates generation - Extracellular simulation* – for further details), and especially on drift (Drifting column). When simulating drifting templates, in fact, the number of actual extracellular spikes for each cell model is N templates times N drift steps. Note that in order to generate the *far-neurons* noise model, the minimum amplitude should be set to 0, so that low-amplitude templates are not discarded. The number of templates available in the template library will be the specified number of templates (N templates) times the number of cell models (13 by default).

**Table 1:** Templates and recordings generation run times depending on several simulation parameters. The templates simulations were run with 13 concurrent jobs (same as the number of models). The recordings simulations were run with four concurrent jobs, and a chunk duration of 20 s.

| Templates generation |  |  |  |  |  |  |
| --- | --- | --- | --- | --- | --- | --- |
| Probe | N templates | N channels | Min. amplitude | Drifting | N drift steps | Run time (s) |
| tetrode-mea-l | 30 | 4 | 30 | No | - | 169 |
| tetrode-mea-l | 100 | 4 | 30 | No | - | 588 |
| tetrode-mea-l | 100 | 4 | 0 | No | - | 236 |
| Neuronexus-32 | 100 | 32 | 30 | No | - | 567 |
| Neuropixels-128 | 100 | 128 | 30 | No | - | 809 |
| SqMEA-10-15 | 30 | 100 | 30 | No | - | 1027 |
| Neuronexus-32 | 30 | 32 | 30 | Yes | 50 | 2000 |
| Recordings generation |  |  |  |  |  |  |
| Probe | N cells | N channels | Duration | Bursting | Drifting | Run time (s) |
| tetrode-mea-l | 6 | 4 | 10 | No | No | 0.5 |
| tetrode-mea-l | 6 | 4 | 600 | No | No | 7 |
| Neuronexus-32 | 20 | 32 | 30 | No | No | 8 |
| Neuronexus-32 | 20 | 32 | 30 | Yes | No | 52 |
| Neuropixels-128 | 60 | 128 | 30 | No | No | 46 |
| SqMEA-10-15 | 50 | 100 | 30 | No | No | 33 |
| Neuronexus-32 | 4 | 32 | 60 | No | Yes | 15 |
| Neuronexus-32 | 20 | 32 | 60 | No | Yes | 37 |

Recordings are then generated using the simulated template libraries. In Table 1, the *Recordings generation* section shows run times for several recordings with different probes, durations, number of cells, bursting, and drifting options. The main parameter that affects simulation times is the number of cells, as it increases the number of modulated convolutions. Bursting and drifting behavior also increase the run time of the simulations, because of the extra processing required in the convolution step. The simulation run times, however, range from a few seconds to a few minutes. Therefore, the speed of `MEArec` enables users to generate numerous recordings with different parameters for testing spike sorter performances. Moreover, the software internally uses memory maps to reduce the RAM usage and the simulations can be *chunked* in time. These features enable users to simulate long recordings on probes with several hundreds of electrodes (e.g. Neuropixels probes) without the need of large-memory nodes or high-performance computing platforms.

## 5 Simulation output

The templates generation outputs a `TemplateGenerator` object, containing the following fields:

`templates` contains the generated templates – array with shape (n_templates, n_electrodes, n_points) for non-drifting templates or (n_templates, n_drift_steps, n_electrodes, n_points) for drifting ones

`locations` contains the 3D soma locations of the templates – array with shape (n_templates, 3) for non-drifting templates or (n_templates, n_drift_steps, 3) for drifting templates.

`rotations` contains the 3D rotations applied to the cell model before computing the template – array with shape (n_templates, 3) (for drifting templates rotation is fixed)

`celltypes` contains the cell types of the generated templates – array of strings with length (n_templates) `info` contains a dictionary with parameters used for the simulation (`params` key) and information about the probe (`electrodes` key)

The recordings generation outputs a `RecordingGenerator` object, containing the following fields:

`recordings` contains the generated recordings – array with shape (n_electrodes, n_samples) `spiketrains` contains the spike trains – list of (n_neurons) `neo.Spiketrain` objects [15]

`templates` contains the selected templates – array with shape (n_neurons, n_jitters, n_electrodes, n_templates samples) templates for non-drifting recordings - or (n_neurons, n_drift_steps, n_jitters, n_electrodes, n_neurons) for drifting ones

`templates_celltypes` contains the cell type of the selected templates – array of strings with length (n_neurons)

`templates_locations` contains the 3D soma locations of the selected templates – array with shape (n_neurons, 3) for non-drifting recordings or (n_neurons, n_drift_steps, 3) for drifting ones

`templates_rotations` contains the 3D rotations applied to the selected templates – array with shape (n_neurons, 3)

`channel_positions` contains the 3D positions of the probe electrodes – array with shape (n_electrodes, 3)

`timestamps` contains the timestamps in seconds – array with length (n_samples)

`voltage_peaks` contains the average voltage peaks of the templates on each electrode – array with shape (n_neurons, n_electrodes)

`spike_traces` contains a clean spike trace for each neuron (generated by a clean convolution between the spike train and the template on the electrode with the largest peak) – array with shape (n_neurons, n_samples)

`info` contains a dictionary with parameters used for the simulation

When simulating with the Python API, the returned `TemplateGenerator` and `RecordingGenerator` can be saved as `.h5` files with:

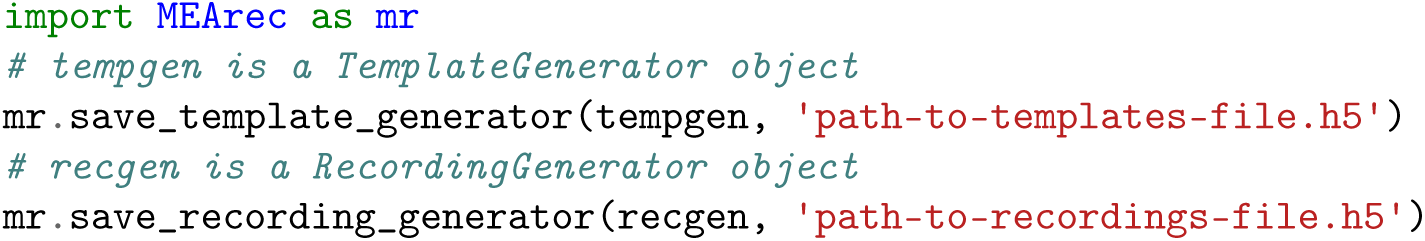

The generation using the CLI saves templates and recordings directly. The saved templates and recordings can be loaded in Python as `TemplateGenerator` and `RecordingGenerator` objects with:

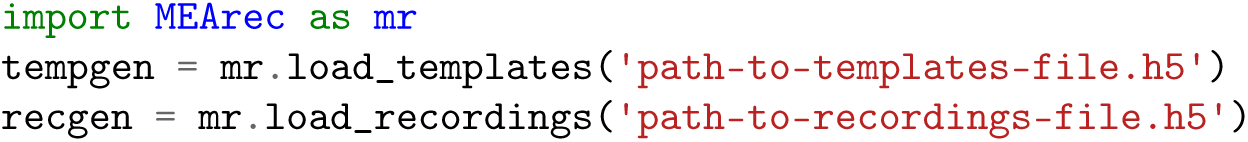

## 6 Discussion

In this paper we have presented `MEArec`, a Python package for simulating extracellular recordings for spike sorting development and validation. We first showed the ease of use of the software, whose command line interface and simple Python API enable users to simulate extracellular recordings with a couple of commands or a few lines of code. We then introduced an overview of the software function, consisting in separating the *templates* and the *recordings* generation to improve efficiency and simulation speed. We explored the capability of reproducing and controlling several aspects of extracellular recordings which can be critical for spike sorting algorithms, including spikes in a burst with varying spike shapes, spatio-temporal overlaps, drifting units, and noise assumptions. We illustrated two examples of using `MEArec`, in combination with `SpikeInterface` [4], as a testbench platform for developing spike sorting algorithms. Finally, we benchmarked the speed performance of `MEArec` (Table 1).

Investigating the validation section of several recently developed spike sorting algorithms [44, 40, 27, 23, 26, 28, 47], it is clear that the neuroscientific community needs a standardized validation framework for spike sorting performance. Some spike sorters are validated using a so called hybrid approach, in which well-identified units from previous experimental recordings are artificially injected in the recordings and used to compute performance metrics [44, 40, 46]. The use of templates extracted from previously sorted datasets poses some questions regarding the accuracy of the initial sorting, as well as the complexity of the well-identified units. Alternatively, other spike sorters are validated on experimental paired ground-truth recordings [9, 47]. While these valuable datasets [20, 22, 37, 33] can certainly provide useful information, the low count of ground-truth units makes the validation incomplete and could result in biases (for example algorithm-specific parameters could be tuned to reach a higher performance for the recorded ground-truth units). A third validation method consist of using simulated ground-truth recordings [12]. While this approach is promising, in combination with experimental paired recordings, the current available simulators [7, 19, 35] present some limitations in terms of biological realism, controllability, speed, and/or ease of use (see Introduction). We therefore introduced `MEArec`, a software package which is computationally efficient, easy to use, highly controllable, and capable of reproducing critical characteristics of extracellular recordings relevant to spike sorting, including bursting modulation, spatio-temporal overlaps, drift of units over time, and various noise profiles.

The capability of `MEArec` to replicate complexities in extracellular recordings which are usually either ignored or not controlled in other simulators, permits the user to include tailored simulations in the spike sorting implementation process, using the simulator as a testbench platform for algorithm development. `MEArec` simulations could not only be used to test the final product, but specific simulations could be used to help implementing algorithms that are able to cope with drifts, bursting, and spatio-temporal overlap, which are regarded as the most complex aspects for spike sorting performance [43, 47].

In `MEArec`, in order to generate extracellular templates, we used a well-established modeling framework for solving the single neuron dynamics [8], and for calculating extracellular fields generated by transmembrane currents [30, 19]. These models have some assumptions that, if warranted, could be addressed with more sophisticated methods, such as finite element methods (FEM). In a recent work [6], we used FEM simulations and showed that the extracellular probes, especially MEAs, affect the amplitude of the recorded signals. While this finding is definitely interesting for accurately modeling and understanding how the extracellular potential is generated and recorded, it is unclear how it would affect the spike sorting performance. Moreover, when modeling signals on MEAs, we used the method of images [36, 6], which models the probe as a infinite insulating plane and better describes the recorded potentials for large MEA probes [6].

Secondly, during templates generation, the neuron models were randomly moved around and rotated with physiologically acceptable values [5]. In this phase, some dendritic trees might unnaturally cross the probes. We decided to not modify the cell models and allow for this behavior for sake of efficiency of the simulator. The modification of the dendritic trees for each extracellular spike generation would in fact be too computationally intense. However, since the templates generation phase is only run once for each probes, in the future we plan to both to include the probe effect in the simulations and to carefully modify the dendritic positions so that they do not cross the probes’ plane.

Another limitation of the proposed modelling approach is in the replication of bursting behavior. We implemented a simplified bursting modulation that attempts to capture the features recorded from extracellular electrodes by modifying the template amplitude and shape depending on the spiking history. However, more advanced aspects of waveform modulation caused by bursting, including morphology-dependent variation of spike shapes, cannot be modelled with the proposed approach, and their replication requires a full multi-compartment simulation [19]. Nevertheless, the suggested simplified model of bursting could be a valuable tool for testing the capability of spike sorters to deal with this phenomenon.

Finally, the current version of `MEArec` only supports cell models from the Neocortical Microcircuit Portal [32, 42], which includes models from juvenile rat somatosensory cortex. The same cell model format is also being used to build a full hippocampus model [34] and other brain regions, and therefore the integration of new models should be straightforward. However, we also provide a mechanism to use custom cell models. For example, cell models from the Allen Brain Institute database [17]^1^, which contains models from mice and humans, can be easily used to simulate templates and recordings, as documented in this notebook: https://github.com/alejoe91/MEArec/blob/master/notebooks/generate_recordings_with_allen_models.ipynb. Other cell models can be used with the same approach.

The use of fully-simulated recordings can raise questions on how well the simulations replicate real extracellular recordings. For example, recordings on freely moving animals present several motion artifacts that are complicated to model and incorporate into simulators. For these reasons, we believe that spike sorting validation cannot be solely limited to simulated recordings. In a recent effort for spike sorting validation, named SpikeForest [31], the authors have gathered more than 650 ground-truth recordings belonging to different categories: paired recordings, simulated synthetic recordings (including `MEArec`-generated datasets), hybrid recordings, and manually sorted data. We think that a systematic benchmark of spike sorting tools will benefit from this larger collection of diverse ground-truth recordings, and in this light, MEArec can provide high-quality simulated datasets to aid this purpose.

In conclusion, we introduced `MEArec`, which is a Python-based simulation framework for extracellular recordings. Thanks to its speed and controllability, we see `MEArec` to aid both the development and validation spike sorting algorithms and to help understanding the limitation of current methods, to improve their performance, and to generate new software tools for the hard and still partially unsolved spike sorting problem.

## Information Sharing Statement

The presented software package is available at https://github.com/alejoe91/MEArec and https://github.com/alejoe91/MEAutility (used for probe handling). The packages are also available on `pypi`: https://pypi.org/project/MEArec/ - https://pypi.org/project/MEAutility/. All the datsets generated for the paper and used to make figures are available on Zenodo at https://doi.org/10.5281/zenodo.3696926, where instruction to generate figures are also provided.

## Acknowledgments

A.P.B. and G.T.E. are part of the Simula-UCSD-University of Oslo Research and PhD training (SU-URPh) program, an international collaboration in computational biology and medicine funded by the Norwegian Ministry of Education and Research. We would also like to thank Samuel Garcia for the help in improving the performance of the simulator. Finally, we would like to thank Kristian Lensjø, Jennifer Hazen, and Mikkel Lepperød for their valuable feedback on the article.

## Appendix A command line interface (CLI)

`MEArec` implements a command line interface (CLI) to make templates and recordings generation easy to use and to allow for scripting. In order to discover the available commands, the user can use the `--help` option:

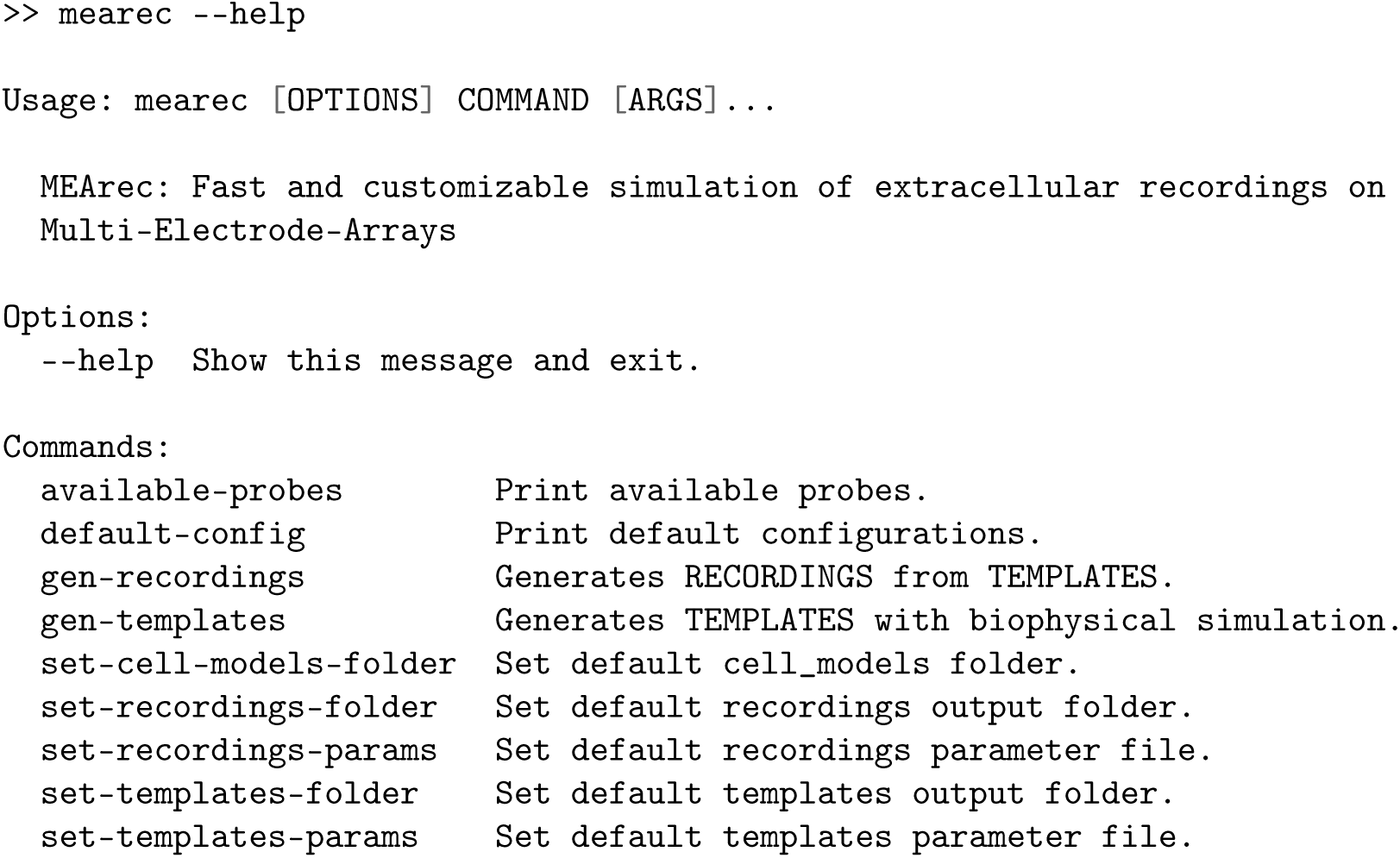

Each available command can be inspected using the `--help` option:

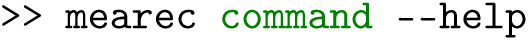

At installation, `MEArec` creates a configuration folder (`.config/mearec`) in which global settings are stored. The default paths to cell models folder, templates and recordings output folders and parameters can be set using the `set-` commands. By default, these files and folders are located in the configuration folder.

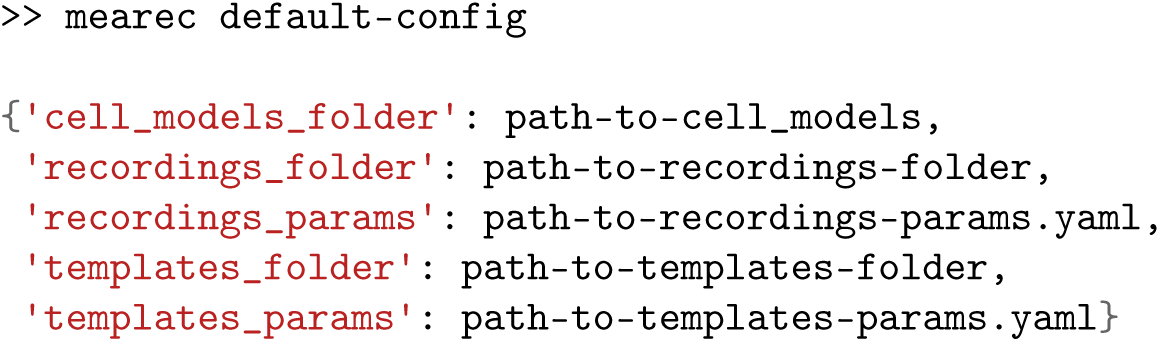

A list of available probes can be found by running the `available-probes` command.

## Supplementary Methods

### Templates generation

This section explains the templates generation phase of the simulator (Figure 1A). Table 2 shows the list of parameters involved in this phase, their default values, types, and an explanation of their function.

**Table 2:** Templates generation parameter list, values, types, and explanations.

| Parameter | Value | Type | Explanation |
| --- | --- | --- | --- |
| **Intracellular simulation settings** |  |  |  |
| sim_time | 1 | float | intracellular simulation time in seconds |
| target_spikes | [3, 50] | list (int) | min-max number of spikes in sim_time |
| cut_out | [2, 5] | list (float) | pre-post peak cut_out in ms |
| dt | 0.03125 | float | time step in ms (default is 32 kHz) |
| delay | 10 | float | stimulation delay in ms |
| weights | [0.25, 1.75] | list (float) | weights to multiply stimulus amplitude if number of spikes is above (0.25) or below (1.25) target spikes |
| **Extracellular simulation settings** |  |  |  |
| rot | physrot | string | rotation to apply to cell models (norot, xrot, yrot, zrot, physrot, 3drot) |
| probe | Neuronexus-32 | string | extracellular probe (if null an available probe is randomly chosen) |
| ncontacts | 10 | int | number of contacts per recording site |
| overhang | 30 | float | extension in $\mu\text{m}$ beyond MEA boundaries for neuron locations (if corresponding lim is null) |
| xlim | [10,80] | list (float) | limits (low, high) for neuron locations in the x-axis in $\mu\text{m}$ |
| ylim | null | list (float) | limits (low, high) for neuron locations in the y-axis in $\mu\text{m}$ |
| zlim | null | list (float) | limits (low, high) for neuron locations in the z-axis in $\mu\text{m}$ |
| min_amp | 30 | float | minimum template amplitude |
| n | 50 | int | number of spikes per cell model |
| n_overlap_pairs | null | int | number of spatially overlapping templates |
| drifting | False | bool | if True, drifting templates are simulated |
| max_drift | 100 | float | maximum distance from the initial and final cell position |
| min_drift | 30 | float | minimum distance from the initial and final cell position |
| drift_steps | 30 | int | number of drift steps |
| drift_xlim | [-10, 10] | list (float) | limits (low, high) for neuron drift locations in the x-axis (depth) |
| drift_ylim | [-10, 10] | list (float) | limits (low, high) for neuron drift locations in the y-axis (depth) |
| drift_zlim | [20, 80] | list (float) | limits (low, high) for neuron drift locations in the z-axis (depth) |
| seed | null | int | random seed for positions and rotations |

`MEArec` is compatible with realistic multi-compartment neuronal models from the Neocortex Microcircuit Portal (NMC - [42, 32]). Upon installation, 13 cell models from layer 5 are copied in the package folder. The set of available neurons can be easily extended with more cell models from the NMC portal. In order to do so, the user can download and unzip the cell models in the cell models folder (which can be retrieved using the `mearec default-config` command). In addition to NMC cell models, `MEArec` also has a custom mechanism to load cell models. For example, the custom mechanism can be used to simulate recordings with models from the Allen Institute of Brain Science [17]^2^, as shown here: https://github.com/alejoe91/MEArec/blob/master/notebooks/generate_recordings_with_allen_models.ipynb.

#### Intracellular simulation

The neuronal model dynamics is solved using the `NEURON` simulator [8]. The neuron’s soma is stimulated with a constant current for a user-defined simulation time (1 secondecond by default - `sim_time` parameter) and the stimulation weight is adjusted (using the `weights` parameter) so that the number of spikes in the simulation period is within a target interval (between 3 and 50 by default - `target_spikes` parameter). The stimulation starts after `delay` ms from the start of the simulation to avoid initialization artifacts. The simulation time step is defined by the parameter `dt` (default is 0.03125 ms, corresponding to 32 kHz). Single spikes are then detected by threshold crossing, aligned, and cropped (using the `cut_out` parameter). The transmembrane currents of all segments are saved to disk, so that the intracellular simulation only needs to be run once for each cell model.

#### Extracellular simulation

Transmembrane currents generated by the *intracellular simulation* are used to compute extracellular potentials at the electrode locations using `LFPy` [18]. Transmembrane currents are distributed over a line source with the length of its corresponding neural segment. Using the quasi-static approximation [38] and with the assumption of a homogeneous, isotropic, and infinite neural tissue with conductivity *σ* = 0.3 S/m [16], the contribution of each compartment *i* at position ***r***_*i*_ with transmembrane current *I*_*i*_(*t*) to the electric potential on an electrode at position ***r***_*j*_ reads [24, 18, 5]:

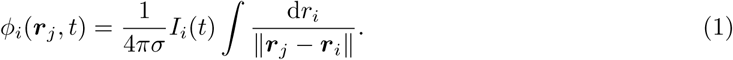

While the assumption of an infinite milieu holds for small probes, such as microwires and tetrodes, when using larger silicon probes, the use of the method of images (MoI) [36] can yield a better estimate of the extracellular potential [6]. Using MoI, the contribution of a transmembrane current to an electrode at position ***r***_*j*_ reads:

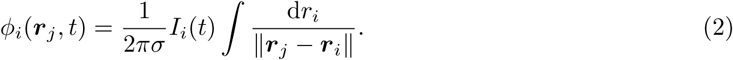

The simulated extracellular spike is obtained by summing up the contributions of all compartments. For each recording site, the electric potential can be computed on several points within the electrode area (`ncontacts` parameter - 10 points by default), that are then averaged to model the spatial filtering properties of the electrodes (disk-electrode approximation [30]).

Each cell model, during the *templates generation* phase, is used to generate several spikes (`n` parameter - 50 by default). For each extracellular action potential, the neuron is randomly moved to a position within user-defined boundaries (`xlim`, `ylim`, `zlim` parameters). If the boundary for a specific axis is set to `null`, the limits are computed as the boundary of the probe in that axis plus the `overhang` value (default 30 µm). Moreover, a random rotation of the model can be optionally added (`rot` parameter). The models can be only shifted (norot), rotated along a single axis (xrot, yrot, zrot), rotated with a physiological rotation (physrot), or rotated randomly along all axes (3drot). For further details we refer to [5]. Extracellular spikes are included in the dataset only if their maximum amplitude is greater than a user-defined minimum amplitude (`min_amp` parameter - 30 µV by default). In order to use the *far-neurons* noise model (Figure 7), the minimum amplitude parameter should be set to 0, so that low amplitude templates are not discarded.

#### Probe models

Probe models are handled using the `MEAutility` Python package (https://github.com/alejoe91/MEAutility), which is automatically installed upon `MEArec` installation. The probe type can be chosen using the `probe` parameter (if not set, a random probe will be selected). `MEAutility` contains a large variety of available probe designs, e.g. commercial Neuronexus probes, Neuropixels [27], and high-density square MEA (Figure 3), and it also allow users to define new probes using a `yaml` file or a Python dictionary. The probe definition contains information about the number and arrangement of the electrodes, the electrode shape and size (used for spatial filtering), the plane in which electrodes are located, and the probe type (`wire` or `mea`), which tells the simulator whether to use the infinite assumption (Equation 1) or MoI (Equation 2) for the extracellular potential calculation. In order to list the available probes and their information, one can use the `mearec available-probes --info` command.

#### Drifting templates

When inserting recording probes in the brain, over time there might be relative movement between the probe and the tissue, which causes a so-called drift in the recorded action potentials. In order to incorporate this phenomenon in the simulation of the recordings, drifting templates has to be generated (when the `drifting` parameter is set to true). From an initial random position of the cell model which satisfies the requirements in terms of location (within boundaries) and amplitude (above the detection threshold) a final drifting position is found so that the same conditions are satisfied. Moreover, the user can choose the preferred drifting direction by setting the `drift_xlim`, `drift_ylim`, and `drift_zlim` parameters, which control the drifting limits from the initial position. The minimum and maximum drifting distance can also be set (with the `min_drift` and `max_drift` parameters). When the final position is selected, the cell model is moved along a straight line connecting the initial and final position and the extracellular spike is simulated for `drift_steps` equidistant points (30 points by default) along this line (Figure 6A).

The templates generation phase can be reproduced by setting the `seed` parameter, which is randomly selected if it is set no `null`.

### Recordings generation

When a template library is generated, it can be used to generate many recordings, as shown in Figure 1B. Tables 3 and 4 show the list of parameters involved in the recordings generation phase, their default values, types, and an explanation of their function.

**Table 3:** Recordings generation parameter list, values, types, and explanations.

| Parameter | Value | Type | Explanation |
| --- | --- | --- | --- |
| Spike trains |  |  |  |
| n_exc | 7 | int | number of excitatory cells |
| n_inh | 3 | int | number of inhibitory cells |
| f_exc | 5 | float | average firing rate of excitatory cells in Hz |
| f_inh | 15 | float | average firing rate of inhibitory cells in Hz |
| st_exc | 1 | float | firing rate standard deviation of excitatory cells in Hz |
| st_inh | 3 | float | firing rate standard deviation of inhibitory cells in Hz |
| min_rate | 0.5 | float | minimum firing rate in Hz |
| ref_per | 2 | float | refractory period in ms |
| process | poisson | string | process for spike train simulation (poisson-gamma) |
| gamma_shape | 2 | float | gamma shape (for gamma process) |
| duration | 10 | float | duration in seconds |
| Cell types |  |  |  |
| excitatory | ['PC', 'SS', 'SP'] | list (string) | Excitatory cell types |
| inhibitory | ['AC', 'BP', 'BC', 'BTC', 'ChC', 'DBC', 'MC', 'NGC'] | list (string) | Inhibitory cell types |
| Templates |  |  |  |
| min_dist | 25 | float | minimum distance between neurons |
| min_amp | 50 | float | minimum spike amplitude in μV |
| max_amp | 500 | float | maximum spike amplitude in μV |
| xlim | null | list (float) | limits for neuron x in μm (min, max) |
| ylim | null | list (float) | limits for neuron y in μm (min, max) |
| zlim | null | list (float) | limits for neuron z in μm (min, max) |
| overlap_threshold | 0.9 | float | threshold to consider two templates spatially overlapping |
| n_jitters | 10 | int | number of temporal jittered copies for each template |
| upsample | 8 | int | upsampling factor to extract jittered copies |
| pad_len | [3, 3] | list (float) | padding of templates in ms |

**Table 4:** Recordings generation parameter list, values, types, and explanations.

| Parameter | Value | Type | Explanation |
| --- | --- | --- | --- |
| **Recordings** |  |  |  |
| fs | null | int | sampling frequency in Hz<br>(if null it is computed from the templates) |
| dtype | float32 | dtype | dtype of generated recordings |
| overlap | False | bool | if True, temporal and spatial overlap are computed for each spike (it may be time consuming) |
| extract_waveforms | False | bool | if True, waveforms are extracted from recordings |
| sync_rate | null | float | synchrony rate ([0-1]) for spike trains of spatially overlapping templates |
| sync_jitt | 1 | float | jitter in ms for added synchronous spikes |
| modulation | electrode | string | type of modulation [none \| template \| electrode]<br>none - no modulation<br>template - each spike instance is modulated with the same value on each electrode<br>electrode - each electrode is modulated separately |
| sdrand | 0.05 | float | standard deviation of Gaussian modulation |
| bursting | False | bool | if True, spikes are modulated in amplitude depending on the ISI |
| exp_decay | 0.1 | float | (bursting) experimental decay in amplitude between consecutive spikes |
| n_burst_spikes | 10 | int | (bursting) max number of 'bursting' consecutive spikes |
| max_burst_duration | 100 | float | (bursting) duration in ms of maximum burst modulation |
| shape_mod | False | bool | if True waveforms are stretched in shape with a sigmoid transform depending on their modulation value |
| shape_stretch | 30 | float | amount of stretch for shape modulation |
| n_bursting | null | int | number of bursting units. If null all units are bursting |
| chunk_duration | 20 | float | chunk duration (if running into MemoryError) |
| noise_level | 10 | float | noise standard deviation in $\mu V$ |
| noise_mode | uncorrelated | string | [uncorrelated \| distance-correlated \| far-neurons] |
| noise_color | False | bool | if True noise is colored resembling experimental noise |
| noise_half_distance | 30 | float | (distance-correlated) distance between electrodes in $\mu m$ for which correlation is 0.5 |
| far_neurons_n | 300 | int | (far-neurons) number of far neurons to be simulated |
| far_neurons_max_amp | 10 | float | (far-neurons) maximum amplitude of far neurons |
| far_neurons_noise_floor | 0.5 | float | (far-neurons) percent of additive random noise |
| far_neurons_exc_inh_ratio | 0.8 | float | (far-neurons) excitatory / inhibitory noisy neurons ratio [0-1] |
| color_peak | 300 | float | (color) peak / cutoff frequency of resonating filter in Hz |
| color_q | 2 | int | (color) quality factor of resonating filter |
| color_noise_floor | 0.5 | float | (color) percent of additive random noise |
| filter | True | bool | if True recordings are filtered |
| filter_cutoff | [300, 6000] | float/list | filter cutoff frequencies in Hz |
| filter_order | 3 | int | filter order |
| drifting | False | bool | if True drifting recordings are simulated |
| n_drifting | null | int | number of drifting units. If null all units are drifting |
| preferred_dir | [0, 0, 1] | list | preferred drifting direction<br>([0, 0, 1] is positive z, direction) |
| angle_tol | 15 | float | tolerance for direction in degrees |
| slow_drift_velocity | 5 | float | slow drift velocity in $\mu m/min$ |
| fast_drift_period | float | 20 | period between fast drift events in s |
| fast_drift_max_jump | float | 20 | maximum amplitude jump in $\mu V$ for fast drifts |
| fast_drift_min_jump | float | 5 | minimum amplitude jump in $\mu V$ for fast drifts |
| t_start_drift | 0 | float | time in seconds after which drift starts |
| **Seeds** |  |  |  |
| spiketrains | int | null | seed for spike trains generation |
| templates | int | null | seed for template selection |
| convolution | int | null | seed for convolution operation |
| noise | int | null | seed for noise generation |

#### Spike trains generations

In order to obtain the spiking activity, spike trains have to be generated. All the spike train generation parameters can be found in the `spiketrains` section of the recordings parameters.

Spike trains can be generated either as Poisson or Gamma processes (`process` parameter). If the Gamma process is selected, its shape is controlled by the `gamma_shape` parameter (default is 2). The user can decide the number of excitatory (`n_exc`) and inhibitory neurons (`n_inh`) in the recordings. The average and standard deviation of the firing rates of excitatory and inhibitory neurons can be chosen (with the `f_exc`, `f_inh`, `st_exc`, and `st_inh` parameters), as well as the minimum accepted firing rate (`min_rate` - default 0.5 Hz). Alternatively, the user can define the type (E-I) and mean firing rate of all neurons in the recordings. As Poisson and Gamma processes do not have a minimum inter-spike-interval, spikes violating a refractory period (`ref_per` - 2 ms by default) are removed from the spike trains. Finally, the duration of the spike trains sets the duration of the recordings (`duration` parameter). Spike trains are represented as `neo.SpikeTrain` objects [15].

#### Excitatory and inhibitory cell types

The `cell_types` section of the recordings parameters tells the simulator which cell types are excitatory and which are inhibitory. For all cell models in the Neocortical Microcircuit Portal [42], excitatory cells can be pyramidal cells (PC), star pyramidal cells (SP), and stellate cells (SS). The population of inhibitory cells is more diverse and it includes: axon cells (AC), bipolar cells (BP), bitufted cells (BTC), basket cells (BC), Chandelier cells (ChC), double bouquet cells (DBC), Martinotti cells (MC), and neurogliaform cells (NGC) [32]. This substrings are used to identify the cell models belonging to the excitatory and inhibitory group for the template selection process. When using custom models, this dictionary should be overwritten for a correct selection of excitatory and inhibitory templates.

#### Template selection and pre-processing

After spike trains are generated, templates are selected from the template library and associated with each spike train. The parameters involved in the template selection and pre-processing are in the `templates` section of the recordings parameters.

Templates are chosen based on amplitude, distance, spatial overlap, and cell type. The selection algorithm discards templates with a peak amplitude below and above user-defined threshold (`min_amp` and `max_amp` parameters) and with a distance from already selected neurons below a minimum distance (`min_dist` parameter). Moreover, the user can select specific boundaries in the x-, y-, and z-direction (`xlim`, `ylim`, and `zlim` parameter). If the boundaries are set to `null` (by default), there is no restriction on the neurons’ location. Templates are chosen so that the number of excitatory and inhibitory types matches the spike trains’ ones. Finally, the user can select the number of spatially overlapping template pairs in the recordings (`n_overlap_pairs` parameter). Two templates A and B are identified as spatially overlapping if the amplitude of template B on the electrode with largest amplitude for template A is above 90% (`overlap_threshold` parameter) of its maximum amplitude, and viceversa.

When templates are selected, they are pre-processed before the convolution operation. First, the templates are padded on both sides (by default extending the templates of 3 ms on each side - `pad_len` parameter) in order to ensure a smooth convolution operation. The template baseline is first removed, then the templates are extended in both directions by linearly interpolating their initial and final values to 0. Finally, this linearly extended template is re-interpolated with a cubic spline.

Next, to model the time variation occurring during sampling, for each template `n_jitter` versions are created (10 by default). Jittering is performed by upsampling the templates (8x by default - `upsample` parameter) and shifting them randomly in time within a sampling period, before downsampling them back to the original sampling frequency.

#### Recordings construction

In the `recordings` section of the recordings parameters, the user can set several parameters for the recordings generation. If not specified, the sampling frequency of the recordings (`fs` parameter) is the same as the generated templates (32 kHz by default), but the user can choose a different sampling rate. In this case the templates are resampled using a polyphase filter. If the `overlap` parameter is set to true, each spike is annotated as `NO` (no overlap), `TO` (temporal overlap), or `STO` (spatio-temporal overlap). If the `extract_waveforms` parameter is set to true, after the recordings generation the waveforms are extracted from the recordings and loaded to the spike train objects. The simulation can also be performed in temporal chunks by setting the `chunk_duration` parameter (20 s by default). Chunking is used to reduce the amount of RAM required by the simulation. Different chunks can also be processed in parallel by providing an `n_jobs` (>1) argument when launching the simulation.

#### Overlapping spikes and spatio-temporal synchrony

Spatio-temporal overlapping of spikes can make spike sorting very challenging [40, 47]. In order to control how spike sorting is affected by the rate of overlapping spikes, `MEArec` enables users to modify the spike trains in order to introduce a controlled amount of spatio-temporal overlapping synchrony (Figure 5).

If the synchrony rate is set (`sync_rate` parameter), the spike trains of spatially overlapping templates are modified to reach the desired synchrony rate. If the chosen synchrony rate is lower than the initial rate, spatio-temporal overlapping spikes are randomly removed from the spike trains. Conversely, when the chosen synchrony rate is greater than the initial rate, additional spikes that do not violate the refractory period are randomly added to the corresponding spike trains until the desired rate is reached. The additive spikes are jittered randomly within a user-defined interval (`sync_jitt` - default ±1 *ms*).

#### Modulated convolution

Pre-processed templates and spike trains are combined with a customized (modulated) convolution. For each spike event of a spike train, a randomized jittered version of the corresponding template is selcted, to reproduce variations due to the finite sampling rate. In order the mimic the variability of spikes in experimental data and computational models [1, 19], the convolution between spike trains and templates is modulated, i.e., the template corresponding to each spike can be modified both in amplitude and in shape (Figure 4).

There are three types of amplitude `modulation` available: 1) *none* (no modulation), 2) *template*, 3) *electrode* modulation (default). On top of amplitude modulation, when `modulation` is not *none*, shape modulation can be used by setting the `shape_mod` parameter to true.

#### Amplitude modulation

The amplitude modulation consists of scaling the amplitude of each spike event with a modulation value. When the *template* modulation is selected, the modulation value is the same for all the electrodes. When the *electrode* modulation is used, each electrode has a slightly different modulation value. For the *template* and *electrode* modulation types, if the `bursting` parameter is set to false, the modulation value is a random value drawn from a normal distribution *𝒩* (1, `sdrand`^2^), (where `sdrand` is 0.05 by default). As the distribution has mean equal 1, the average amplitude of the resulting modulated spikes is the same as the original template. When the `bursting` parameter is set to true, the modulation values are computed to reproduce the amplitude scaling due to bursting behavior (see Figure 4A). The user can choose how many units will be affected by bursting (`n_bursting` parameter). Consecutive spikes occurring within a user-defined bursting period (`max_burst_duration` parameter - default 100 ms) are scaled with a sub-linear function (up to a maximum number of consecutive spikes `n_burst_spikes` - 10 by default). The amplitude scaling for the *i*-th consecutive spike within a bursting event is computed as:

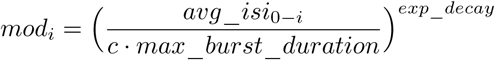

where *avg*_*isi*_0−*i*_ is the average inter-spike-interval (ISI) from the first bursting spike to the current spike in the bursting event, *c* is the number of consecutive spikes encountered up to spike *i*, `max_burst_duration` is the maximum bursting period (default 100 ms), and `exp_decay` is the exponent (0.1 by default). Additionally, the ISI-dependent modulation value is scaled a by a random value drawn from a normal distribution at the template level (*template* modulation) or electrode level (*electrode* modulation).

#### Shape modulation

When `shape_mod` is set to true, spikes are also modulated in shape. Shape modulation consists of stretching the template depending on its modulation value (the same modulation value is used both for amplitude and shape modulation). The stretch is achieved in the following way: first, the template time axis is centered to the template peak and scaled so that its length is equal to 1 – we will refer to this centered and normalized time axis as *x*_*c*_; second, *x*_*c*_ is multiplied by the `shape_stretch` parameter, which controls the amount of stretch – we will refer to this transformed time axis as *x*_*t*_; then, a stretch factor *s* is computed for the entire template (the same factor is computed for all electrodes) as the average modulation value of all electrodes (if *electrode* modulation is used); if the stretch factor is less than 1, *x*_*t*_ is projected on a sigmoid function:

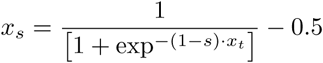

*x*_*s*_ is re-scaled so that its length is `shape_stretch`, and it is now a non-linear stretched time axis. The template is interpolated on *x*_*s*_ with a cubic spline and transformed back to a linear time axis *x*_*r*_ and scaled in amplitude using the modulation value. Figure S1 shows the different axes involved in shape modulation, and examples of the template transformation at different stages for two modulation values (0.9 and 0.7).

#### Noise models and post-processing

Additive noise is superimposed to the signals after the modulated convolution is finished. There are three types of noise models that can be set using the `noise_mode` parameter: *uncorrelated, distance-correlated*, and *far-neurons*. The uncorrelated noise model is an additive Gaussian noise with a user-defined standard deviation (`noise_level` parameter - 10 µV by default). The distance-correlated mode generates a multivariate normal noise with a covariance matrix dependent on the distance between electrodes. The covariance between electrode *i* and *j* is defined as *c*_*ij*_ = *dh/*2*·d*_*ij*_, where *d*_*ij*_ is the distance between the electrodes and *d*_*h*_ is the distance at which the covariance is 0.5 (`noise_half_distance` parameter - 30 µm by default). Finally, the *far-neurons* model generates noise as the activity of many neurons (`far_neurons_n` parameter - 300 by default) with small amplitudes (below `far_neurons_max_amp` - 10 µV by default). The population of distant neurons has an excitatory/inhibitory ratio of `far_neurons_exc_inh_ratio` (default 0.8). A random noise floor with a standard deviation of `far_neurons_noise_floor` (default 0.5) times the standard deviation of the distant neurons’ spiking activity is added, in agreement with experimental data [7].

**Figure S1:**
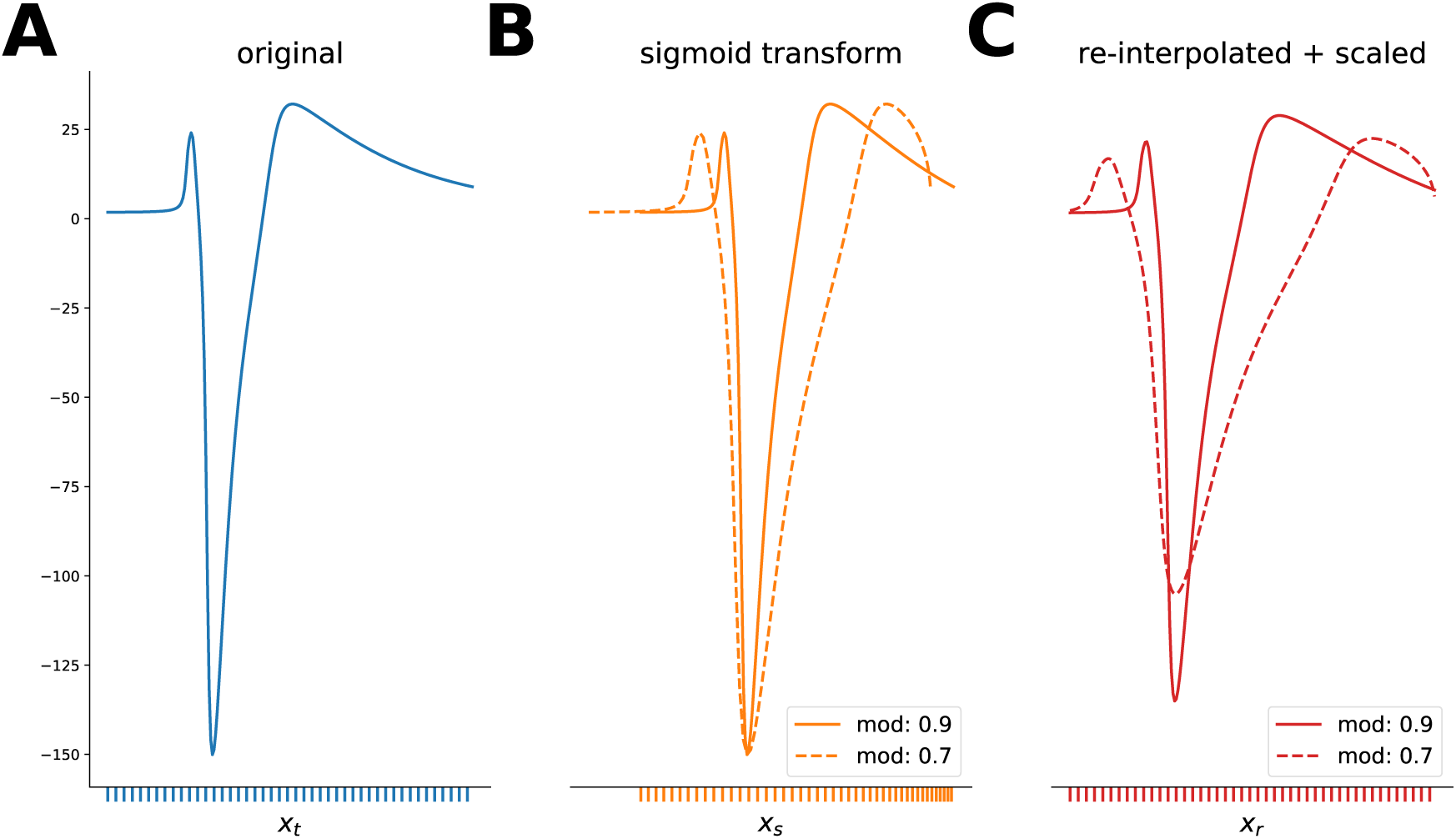
Shape modulation. (A) Original template on a the linear axis *x*_*t*_. (B) Templates after projection on the sigmoid-transformed axis *x*_*s*_ (solid: modulation value=0.9, dashed: modulation value=0.7). Note that different modulation values, and in turn different *s* values, make the *x*_*s*_ range vary. (C) Templares after re-interpolation on the linear *x*_*r*_ and amplitude scaling by the modulation value (solid: modulation value=0.9, dashed: modulation value=0.7).

Uncorrelated and distance-correlated noise types can also be modulated in frequency to match the spectrum observed in experimental data [7, 18]. Extracellular spiking activity exhibit a peak in frequency at around 300 Hz, a 1*/f* spectrum, and a random noise floor. Noise can be *colored* (when the `noise_color` parameter is true) with a second order infinite impulse response (IIR) peak filter and an additional gaussian noise floor. The frequency peak, quality factor, and weight of the random noise floor can be set with the `color_peak`, `color_q`, and `color_noise_floor` parameters. Note that with distance-correlated noise the correlation is slightly reduced by the color filter, as a random noise floor is added.

Optionally, the signals can be filtered (by setting the `filter` to true) with an high-pass or band-pass Butterworth filter of order `filter_order` (3 by default) and cutoff frequencies of `filter_cutoff` ([300, 6000] Hz by default).

#### Drifting recordings

When the `drifting` parameter is set to true, drifting recordings are generated. The template library must have been generated with the `drifting` mode as well. The user can decide the number of drifting units (`n_drifting` parameter). If `n_drifting` is `null`, all units will be drifting.

The generation of drifting recordings is only different in the template selection and modulated convolution steps. In the template selection, in addition to the selection rules based on template amplitude, inter-neuron distance, and spatial overlap, templates are selected if the angle between the drifting direction (computed as the vector connecting the final and initial position) and a user-defined preferred direction (`preferred_dir` parameter - [0, 0, 1] by default) is within an angle tolerance (`angle_tol` parameter - 15° by default).

There are three types of drift modes available (`drift_mode` parameter): *slow, fast*, and *slow+fast*. The different modalities vary in terms of how the drifting template is selected for each spike during the modulated convolution.

For *slow* drifts, a new position is calculated moving from the initial position along the drifting direction with a velocity of `slow_drift_velocity` (default 5 µm*/min*). If a boundary position is reached (initial or final positions), the drift direction is reversed.

For *fast* drifts, the user can set the frequency at which fast drift events occur (every `fast_drift_period` s). When a fast drift event happens, a new template position is selected randomly among the drifting templates for each drifting neuron. The amount of *jump* is controlled by the `fast_drift_min_jump` and `fast_drift_max_jump` in the following way. Let us call *T*_*old*_ the drifting template before the drift event, and *T*_*rand*_ the randomly selected template, which is a candidate for the new template after the fast drift event. *T*_*rand*_ is accepted as the new template only if the difference in amplitude *diff*_*amp*_ between *T*_*old*_ and *T*_*rand*_ on the channel in which *T*_*old*_ has the largest peak is `fast_drift_min_jump` < *diff*_*amp*_ < `fast_drift_max_jump`. This is to ensure that fast drifts are not too abrupt.

Finally, when the *slow+fast* mode is selected, the two previously described modes are combined. In all cases, the user can decide to start the drift `t_start_drift` seconds after the start of the recordings.

#### Reproducibility and seeds

In order to ensure full reproducibility of the recordings, the `seeds` section of the recordings parameters enables users to set several random seeds involved in the simulations. The `spiketrains` seed controls the random generation of spike trains, the `templates` seed the selection of templates from the template library, the `convolution` seed is for all the processes involved in the convolution phase (modulation, jittering, drifting), and the `noise` seed controls the randomness in the noise generation. If any of the seed is not set, a random seed is generated and saved in the recording output (in the `info` dictionary), so that the recordings could be reproduced in the future.

## Footnotes

1 https://celltypes.brain-map.org/

2 https://celltypes.brain-map.org/

